# Time-averaged and Time-varying Structure of the Gastric Network Revealed Through fMRI-Electrogastrogram Synchronization

**DOI:** 10.64898/2026.08.26.747287

**Authors:** Ye’ela Zair, Galia Avidan

**Author notes:** Correspondence should be addressed to: Galia Avidan.

## Abstract

The gastric network, comprised of brain regions whose activity synchronizes with the stomach’s slow-wave rhythm, offers a unique window into the brain-body interaction involved in interoceptive processing. While previous work has established the existence of this network, its intrinsic organization and temporal unfolding remain poorly understood. Here, we reanalyzed resting-state fMRI-electrogastrogram data from 43 healthy adults of both sexes to characterize the time-averaged architecture and time-varying reconfiguration of the gastric network. We identified regions exhibiting phase-locked synchronization with the stomach slow electrical rhythm (0.05 Hz) and characterized cortical parcels comprising this network. Time-averaged graph-theoretical analysis revealed a fixed unimodal organization of functional communities, with primary visual, default mode network (DMN) and dorsal attention regions emerging as the principal time-averaged hubs. Next, we applied edge-centric functional connectivity (eFC) to capture the network state during transient high-amplitude “bursts”. Time-varying community detection revealed communities whose compositions formed integrative combinations of DMN, visual, attentional and control elements. Edge-derived hubs shifted away from primary visual dominancy in the time-averaged analysis, and were instead directed by DMN regions, suggesting that moments of heightened connectivity in the network are coordinated by multisensory integration rather than passive sensory processing. These findings demonstrate that the gastric network is not merely a time-averaged, sensory-bound system, but rather a flexible and dynamically reconfiguring interoceptive network whose organization is selectively coordinated by transient cofluctuation events. This work provides a comprehensive network analysis of gastric-brain coupling and reveals a temporally structured mode of interoceptive integration that may support adaptive physiological and cognitive regulation.

## 1. Introduction

Interoception refers to the sensing, interpretation, and regulation of internal bodily states (Craig, 2003; Khalsa et al., 2009; Chen et al., 2020). Through bidirectional brain-body communication, visceral signals inform emotional regulation, homeostasis, and motivated behavior, for example feeding (Tsakiris and Critchley, 2016; Critchley and Garfinkel, 2017; Tallon-Baudry, 2023). A central component of this communication is the brain-gut axis. In the stomach, interstitial cells of Cajal generate an intrinsic slow-wave rhythm with a cycle of approximately 20 s, which can be measured noninvasively using electrogastrography (EGG) (Martchenko et al., 2020). Recent fMRI-EGG studies have shown that spontaneous BOLD activity in sensory, motor, cingulate, and associative regions synchronizes with this rhythm, forming the “gastric network” (Rebollo et al., 2018; Levakov et al., 2023). This network provides a unique model for examining how slow visceral rhythms are embedded within large-scale cortical organization.

Gastric-brain communication is supported by both anatomical and oscillatory pathways. The vagus nerve provides a major route for bidirectional signaling, conveying sensory information from the stomach to the brain and modulating gastric motility through efferent control (Folgueira et al., 2014; Fülling et al., 2019). Vagal afferents relay visceral information through brainstem and forebrain systems involved in allostatic, emotional, and motivational regulation (Teckentrup and Kroemer, 2023). In parallel, the stomach’s slow rhythm aligns with cortical activity; Gastric phase has been shown to modulate cortical alpha power, with coupling localized to regions including the parieto-occipital sulcus and anterior insula, suggesting that visceral rhythms can shape cortical dynamics (Richter et al., 2017; Azzalini et al., 2019). The cortical territories engaged by gastric coupling overlap with interoceptive, somatosensory, attentional, and body-representation systems (Khalsa et al., 2009; Rebollo and Tallon-Baudry, 2022). Recent evidence linking gastric-brain synchronization to mental health further supports the functional relevance of this coupling (Banellis et al., 2025).

Although previous work has mapped the regions synchronized with gastric activity, the intrinsic organization of the gastric network remains poorly understood. Most existing descriptions focus on where gastric coupling occurs but less is known about how these regions interact as a network, which nodes coordinate information flow, and whether the network expresses distinct operational modes. This distinction is important because interoceptive processing is inherently integrative and time-varying: visceral signals must be continuously combined with additional ongoing context to support adaptive regulation (Chen et al., 2020). Thus, characterizing the gastric network requires methods that capture both its stable architecture and its moment-to-moment reconfiguration.

Edge-centric functional connectivity (eFC) provides such a framework. Unlike conventional node-centric functional connectivity, which summarizes the average correlation between brain regions, eFC decomposes functional connectivity into edge time series that track moment-to-moment fluctuations in pairwise coactivity (Faskowitz et al., 2020). This approach enables the identification of transient high-amplitude cofluctuation events, or “bursts,” and reveals overlapping edge communities that may be obscured in time-averaged connectivity matrices (Bassett et al., 2011; Betzel et al., 2023). Applying this framework to the gastric network can therefore reveal whether gastric-synchronized regions remain organized according to canonical resting-state modules or transiently reconfigure into more integrated states.

Here, we reanalyzed a previously published resting-state fMRI – EGG dataset (Levakov, Ganor, et al., 2023) to characterize the time-averaged and time-varying organization of the human gastric network. First, we validated gastric-brain synchrony and used the resulting regions to define the gastric-network nodes. We then examined the network’s time-averaged functional architecture, community structure, and hub organization. Finally, we applied eFC to identify burst-related time-varying communities and hub reconfiguration. This approach allowed us to test whether the gastric network functions as a fixed sensory-anchored system or as a flexible interoceptive network whose organization changes during transient states of heightened cofluctuation.

## 2. Materials and Methods

This study reanalyzes previously published resting-state functional magnetic resonance imaging (fMRI) and electrogastrography (EGG) data (Levakov et al., 2023a) to reveal novel aspects of the gastric network’s time-averaged and time-varying functional connectivity. This study was conducted in accordance with the Declaration of Helsinki and was approved by the Human Subjects Research Committee at Soroka University Medical Center (SUMC), Beer-Sheva, Israel. All participants provided written informed consent, and their privacy rights were observed throughout. Information regarding code accessibility is detailed in the "Data Availability" section.

### 2.1. Participants

The final reanalyzed dataset comprised 43 healthy adult participants (27 female, 16 male; mean age: 25.19 ± 3.35 years), contributing a total of 84 imaging runs. Inclusion criteria required high-quality Electrogastrogram (EGG) signals, a body mass index (BMI) between 18 and 25, no history of gastric disorders, and compliance with MRI safety guidelines. Sixteen of the 59 participants who initially took part in the study were excluded due to poor EGG signal quality or excessive head motion.

Participants were asked to fast for at least 3 hours and to refrain from drinking for at least thirty minutes before scanning to minimize physiological artifacts. Each session included two 15-minute (N=39) or three 10-minute (N=4) consecutive resting-state fMRI runs with simultaneous EGG recording.

### 2.2. MRI Acquisition

MRI data were acquired using a 3T Philips Ingenia scanner (Philips Healthcare, Amsterdam, The Netherlands) with a 32-channel head coil. Each session included a high-resolution T1-weighted anatomical scan, followed by concurrent EGG and rs-fMRI. Fifty four runs were acquired as two 15-minute sessions, and 5 runs as three 10-minute sessions. Participants were instructed to remain awake, still, and fixate on a blank screen.

Functional MRI data used a gradient-echo Echo-Planar Imaging (EPI) sequence with parallel acquisition (SENSE: factor 2.4). Whole-brain coverage (44 transverse slices, 3 mm thickness, no gap; 3x3x3 mm³ voxel size; FOV: 192x192 mm²). Acquisition parameters: TR=2000 ms, TE=25 ms, flip angle=77°, matrix size=64x64. High-resolution T1-weighted anatomical volumes used a 3D turbo field echo (TFE) pulse sequence (1x1x1 mm³ isotropic voxel size, 150 slices).

### 2.3. Electrogastrogram (EGG) Acquisition

EGG recordings were obtained inside the MRI scanner using an MRI-compatible BIOPAC amplifier (EGG-100C, BIOPAC Systems Inc., Goleta, CA, USA). Participants were supine and instructed to minimize motion. Skin was cleaned, and conductive gel applied before attaching standard cutaneous electrodes (EL503, EL508, BIOPAC Systems Inc.) to the abdomen (Rebollo et al., 2018; Wolpert et al., 2020; Levakov, Ganor, et al., 2023). Four electrode pairs recorded independent EGG channels, with a ground electrode on the hip. EGG signals were recorded at 5000 Hz, gain 5000, with a hardware bandpass filter of 0.005–1 Hz to isolate gastric slow-wave activity.

### 2.4. MRI Preprocessing

MRI preprocessing was identical to Levakov et al. (2023a), using fMRIPrep 20.0.6 (Esteban et al., 2019; RRID: SCR_016216), built on Nipype 1.4.2 (Gorgolewski et al., 2011; RRID: SCR_002502). Key steps included: T1w intensity nonuniformity correction (N4BiasFieldCorrection, ANTs 2.2.0; RRID: SCR_004747), skull stripping (antsBrainExtraction.sh, ANTs), tissue segmentation (FAST, FSL 5.0.9; RRID: SCR_002823), and surface reconstruction (recon-all, FreeSurfer 6.0.1; RRID: SCR_001847). Spatial normalization to MNI152NLin6Asym space (Evans et al., 2012) used nonlinear registration (antsRegistration, ANTs).

For BOLD fMRI, a reference volume was coregistered to T1w using bbregister (FreeSurfer). Head motion parameters were estimated with mcflirt (FSL 5.0.9). Confound regression removed framewise displacement (FD) (Power et al., 2014), 6 motion parameters (and their derivatives/squared terms), and 6 principal components from CSF and WM (aCompCor) (Behzadi et al., 2007). Global signal regression was not applied. One key deviation from Levakov et al. (2023a) that was performed in the current study was the application of 3 mm FWHM Gaussian spatial smoothing. This step allowed us to increase the signal to noise ratio and reduce high frequency noise while preserving broader spatial patterns of brain activity.

### 2.5. EGG Preprocessing

EGG preprocessing used custom Python scripts (Levakov et al., 2023a) based on established methods (Rebollo et al., 2018), aiming to enhance signal-to-noise ratio. Raw EGG was initially downsampled from 5000 Hz to 10 Hz to ensure reliable spectral estimation of slow frequencies and subsequently reduced from 10 Hz to 0.5 Hz to match the temporal resolution of fMRI data.

The optimal channel was selected based on maximal spectral power (Welch’s method, 200s window, 150s overlap) within the 0.033–0.066 Hz gastric range and further validated through manual inspection of the chosen dominant channel by the experimenters. This selected channel was bandpass filtered around its peak gastric frequency (±0.015 Hz) using an FIR filter in MNE-Python (RRID: SCR_017042). Signal reliability was assessed based on distinct spectral peak presence, consistency across channels, and stability across runs (Wolpert et al., 2020).

### 2.6. Experimental Design and Statistical Analyses

Statistical analyses were performed for three objectives: 1. Validating brain-gastric synchrony as described in (Levakov et al., 2023a), 2. Characterizing time-averaged functional connectivity, and 3. Characterizing time-varying edge-centric connectivity. Analyses primarily utilized custom Python scripts built upon standard scientific computing libraries and specialized neuroimaging toolboxes.

#### 2.6.1. Brain-Gastric Synchrony

Synchronization was quantified using the phase-locking value (PLV) (Lachaux et al., 1999), measuring consistent phase lag between signals (0-1), ideal for continuous, rhythmic interactions. The first 15 volumes (30s) of both BOLD and EGG time series were excluded. The Hilbert transformation was then applied to extract the instantaneous phase of both signals, from which the phase angle was computed. The absolute value of the mean phase angle difference across time was then computed according to the following equation:

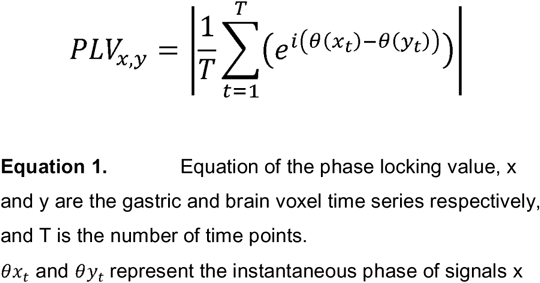

To assess chance-level synchrony, a null model was generated by circularly shifting the gastric signal in time (Harris et al., 2020), ensuring ±60s shifts to break true phase relationships while preserving spectral properties. For 15-min runs, 360 null signals were generated; for 10-min runs, 210. PLV for each voxel was computed against these null signals, and the median null PLV was extracted as a baseline. The subject-level PLV-delta was the difference between empirical and median null PLV, averaged across runs for robustness.

For group-level analysis, individual PLV-delta maps were averaged. A paired t-test between empirical and null PLV maps identified regions with significantly greater gastric synchronization. Multiple comparisons were controlled using a nonparametric permutation test via FSL’s randomize (Winkler et al., 2014; RRID: SCR_006977). A cluster-forming threshold of t=2.3 was applied, with 10,000 permutations. Cluster significance was determined by comparing cluster mass (sum of t-values) to the permutation-based null distribution, with p < 0.05 (FWE-corrected) considered significant.

#### 2.6.2. Voxel-to-ROI Transformation

To define gastric regions of interest (ROIs), significant voxels identified in the group-level brain-gastric synchrony analysis were assigned to predefined cortical parcels using the Schaefer 400-parcel atlas (Schaefer et al., 2018). This atlas provides a fine-grained functional parcellation of the cortex, allowing for anatomically and functionally based ROIs definition. Additionally, the Yeo 17-network parcellation (Yeo et al., 2011) embedded within the Schaefer atlas was used to categorize these ROIs into large-scale resting state cortical networks, facilitating subsequent network-level analyses. Importantly, the Yeo 17-network parcellation was used to gain finer resolution, whereas subdivisions were subsequently summed back to the original Yeo 7-network parcellation for clearer visualization. Each ROI was defined as the intersection between a significant voxel cluster and the corresponding pre-existing cortical parcel (Schaefer et al., 2018). Based on established anatomical characterization methods (Rebollo and Tallon-Baudry, 2022), we have calculated effect size across all significant gastric voxels, by computing Cohen’s d on the difference between empirical and chance level PLV:

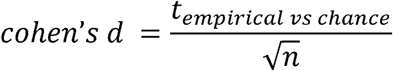

where t_empirical vs chance_ is the t-statistic from the paired t-test between empirical and chance level PLV maps, and n is the number of participants. To obtain the anatomical regions comprising the gastric network, we used the Schafer 2mm cortical parcellation atlas (Schaefer et al., 2018) which contains 200 areas per hemisphere. For each region of the parcellation, we computed the overlap between the gastric network voxels (calculated as the fraction between overlapping voxels out of all parcel’s voxels), alongside averaged effect size across these overlapping gastric network parcels. After calculating the effect size for each parcel, we computed the median Cohen’s d across all gastric networks’ parcels, to obtain a statistically validated cutoff for the regions with the largest effect sizes across hemispheres. Subcortical regions were also included in the initial analysis using the Harvard-Oxford Subcortical Structural Atlas (distributed with FSL; e.g., *FMRIB Software Library*, Smith et al., 2004; Jenkinson et al., 2012) and a probabilistic cerebellar atlas in MNI152 space (Diedrichsen et al., 2009; 2 mm resolution, threshold = 0). After parcellation and selection based on Cohen’s *d* effect size, none of the subcortical or cerebellar regions exceeded the median effect size (Md = 0.426) and were therefore excluded from subsequent network analyses. To obtain stable covariance estimates for computing parcel-wise functional connectivity, we applied the Ledoit-Wolf shrinkage estimator (Ledoit and Wolf, 2003) as implemented in scikit-learn (LedoitWolf(); sklearn.covariance). Ledoit-Wolf shrinkage estimates the covariance as:

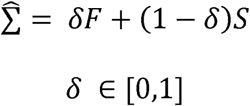

where S denote the empirical covariance matrix and F is the constant-correlation target.

Briefly, the method computes a combination of the empirical covariance matrix and a structured target, using an optimal shrinkage constant, which minimizes the expected distance between the shrinkage estimator and the true covariance matrix. This tends to pull the most extreme coefficients towards more central values, thereby systematically reducing estimation error. This approach reduces noise-driven inflation of extreme covariance values and yields a covariance matrix suitable for downstream correlation (Ledoit and Wolf, 2004). For each subject and run, we fitted the Ledoit-Wolf estimator to the (T × N) parcel time-series matrix and converted the shrunken covariance to a correlation matrix before Fisher transformation.

#### 2.6.3. Time-averaged Functional Connectivity Analysis of the Gastric Network

##### 2.6.3.1. Constructing the Time-averaged Correlation Matrix

Nodes were defined as the brain regions identified with significant gastric-brain synchrony. For each participant, node time-series were the mean BOLD signal across voxels within each ROI. These series were bandpass filtered (0.01–0.1 Hz) to isolate relevant resting-state oscillations (Glerean et al., 2012; Power et al., 2013) and z-scored within participants for amplitude standardization and inter-subject comparison (Betzel et al., 2023; Faskowitz et al., 2021). A 40 × 40 time-averaged FC matrix was constructed for each participant by computing the Pearson correlation between all node time-series, reflecting pairwise functional interactions. Group-level FC was obtained by averaging individual matrices. To assess whether time-averaged functional connectivity exceeded expectations based on intrinsic temporal structure, we generated phase-randomized surrogate time series preserving each node’s power spectrum while disrupting inter-regional synchrony. Functional connectivity matrices were recomputed for each surrogate and compared to empirical estimates.

##### 2.6.3.2. Community structure

To characterize the topological organization of functional connectivity within the gastric network, we applied graph-theoretical measures that quantify features of its modular organization and intrinsic architecture at rest. Modularity represents a fundamental property of both anatomical and functional brain networks, reflecting the balance between functional specialization within local subsystems and the integration of information across them. To infer the community structure of the gastric network, we employed a standard modularity-maximization framework, which partitions network nodes into non-overlapping functional communities. Specifically, the Louvain algorithm (Blondel et al., 2008) was applied to the group-averaged connectivity matrix to identify coherent functional modules (Cho et al., 2023). Owing to the stochastic nature of the algorithm, the partition yielding the maximum modularity index (Q) across 100 iterations was selected as the optimal solution (Sobolevsky and Belyi, 2022). To ensure interpretability, we excluded small communities comprising fewer than three nodes, thereby focusing on the principal functional modules of the gastric network.

##### 2.6.3.3. Centrality measures

To identify hub regions within the gastric network, we quantified several graph-theoretical measures that capture distinct aspects of nodal centrality. We first computed the weighted degree of each node, defined as the sum of connection weights linking that node to all others, such that stronger functional connections contribute proportionally more to nodal degree (Sporns et al., 2007). Nodes exceeding established threshold – weighted degree > 1 SD above the network mean (Sporns et al., 2007), were designated as hubs. To further characterize the functional role of these hubs, we computed each node’s participation coefficient, which quantifies the extent to which a node distributes its connections across multiple communities (Sporns et al., 2007; Van Den Heuvel and Sporns, 2013). Hubs with participation coefficient > 0.3 were classified as connector hubs, reflecting nodes that link multiple functional modules by distributing their connectivity broadly across the network (Tian et al., 2024). In contrast, hubs with participation coefficient < 0.3 were classified as provincial hubs, indicating nodes whose connectivity is concentrated primarily within their own module and which serve as local centers of intra-modular communication (Tian et al., 2024).

#### 2.6.4. Time-varying Analysis of the Gastric Network

##### 2.6.4.1. Extraction of Edge Time Series (eTS)

Node activity time series derived from fMRI data were standardized (z-scored across time points for each node independently). Then, the instantaneous cofluctuation time series for each edge (x,y), (x,y) at each time point t was computed by multiplying the corresponding z-scored signals of the node pair, resulting in an edge time series (eTS) defined as r_{xy}(t)_ - z_x(t)_ z_y(t)_ . This series provides temporally detailed information about the functional connectivity (FC) between edges {x,y} over time, reflecting time-varying functional connectivity (tvFC). Collectively, this procedure produced a cofluctuation matrix E of dimensions T×N_edges_ where T represents the number of fMRI time points and N_edges_ is the total number of node pairs. Thus, each element E_t,e_ corresponds explicitly to the instantaneous cofluctuation value for a specific edge e, at time t. Each column in this matrix represents cofluctuation values for all edges at specific time points, enabling an edge-centric network analysis analogous to traditional node-centric FC assessments. By comparing time series such as r_xy_ - [r_xy_(1),…r_xy_(T)] and r_uv_ - [r_uv_(1),…r_uv_(T)], their similarity was evaluated, producing a vector of coefficients (1×N_edges_) that can be reorganized into a N_edges_ ×N_edges_ correlation matrix, thus representing the similarity in connectivity dynamics between edges. Analyses were conducted independently for each run and each subject. Individual correlation matrices were subsequently averaged across runs and subjects for group-level interpretation.

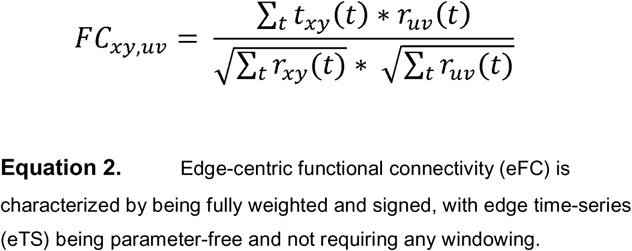

##### 2.6.4.2. Burst Network Construction

Edge time-series (eTS) exhibit ‘burstiness’, i.e., short, high-amplitude cofluctuations reflecting significant, brain-wide network events (Betzel et al., 2023). To quantify global cofluctuation amplitude, we computed the root sum square (RSS) of the edge time series at each time point, yielding an amplitude measure A(t) that captures the magnitude of collective cofluctuations across all edges.

As resting-state data lacks predefined events, burst detection was performed post hoc. To assess whether high-amplitude events reflected genuine network dynamics rather than chance fluctuations, we generated a surrogate distribution by circularly shifting node time series relative to one another, thereby preserving their autocorrelation structure while disrupting cross-node temporal alignment. For each run, 1,000 surrogate realizations were generated. Event frames were identified as time points in which the observed cofluctuation amplitude exceeded the 95th percentile of the surrogate distribution (α = 0.05). In addition, a top-percentile criterion (top 5% of empirical amplitudes) was applied, yielding a hybrid definition of burst events that ensures robustness to both statistical and distributional biases. Burst detection was performed independently for each participant and each run. This procedure yielded approximately 23 burst frames for 15-minute runs and 15 burst frames for 10-minute runs. To assess the robustness of the burst detection procedure, we repeated the analysis using more liberal thresholds, defining bursts as the top 10%, 15%, and 25% of frames based on cofluctuation amplitude. This yielded progressively larger numbers of burst frames across scan durations. Importantly, network analyses produced highly similar results across thresholds, indicating that the main findings are not dependent on the specific choice of threshold. A burst-based adjacency matrix was computed by averaging cofluctuations exclusively from burst frames. This matrix represents a network structure specific to these high-amplitude states. Burst events are expected to show higher correlation compared to time-averaged functional connectivity, and an overlapping architecture, especially in sensorimotor and attentional systems (Levakov et al., 2023b; Faskowitz et al., 2020).

##### 2.6.4.3. Phase-randomization Control Analysis

To assess whether time-varying edge-centric functional connectivity reflected meaningful coordination beyond intrinsic temporal structure, we generated surrogate datasets using phase randomization. For each node time series, we applied a Fourier transform, preserved the amplitude spectrum, and uniformly randomized each edge’s phase components in the range [0, 2π] across all frequency bins, before reconstructing the signal via inverse transform. This procedure preserves each region’s autocorrelation structure while disrupting inter-regional temporal alignment. Surrogate datasets were processed identically to the empirical data, including burst detection and construction of event-based edge-centric functional connectivity (eFC) matrices. Observed EFC patterns were then compared against surrogate-derived distributions to quantify deviations from the null model. Importantly, event detection in surrogate data was performed using identical criteria, ensuring that any observed differences in EFC structure were not attributable to differences in event selection.

##### 2.6.4.4. Edge Community structure

To identify functionally cohesive subnetworks of edges, the Louvain algorithm (Blondel et al., 2008), optimized for edge-based networks (Faskowitz et al., 2021) was used to partition edges into communities. To gain stable community partition, we used the same modularity maximization procedure as in the time-averaged analysis. Following the application of community detection, small edge communities containing less than 10 edges were filtered out to focus on major functional communities of the network. Unlike node-based methods, edge communities allow for overlapping nodal communities when projected onto nodes. This implies that a node can simultaneously participate in multiple edge communities, reflecting its involvement in diverse transient connectivity patterns and providing a nuanced view of dynamic network roles (Betzel et al., 2023; Faskowitz et al., 2020). To quantify the stability of the edge communities, we computed an edge-by-edge co-assignment matrix A. For a set of *P* partitions, we define a matrix *A* of size E×E whose entry A_ij_ is the fraction of partitions in which edges i and j were assigned to the same community:

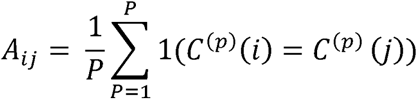

where c^(p)^ (i) denotes the community label of edge i in partition p. The resulting matrix is symmetric, with values ranging from 0 (never co-assigned) to 1 (always co-assigned), and serves as a visualization of the stability and separability of the edge-community structure.

##### 2.6.4.5. Centrality measures

Time-varying hub identification was performed in edge-space by treating each functional connection as a node in the edge-centric correlation graph derived from eTS similarity. Hub detection followed the same criteria as the time-averaged node-based analysis, with weighted degree and participation coefficient computed on the edge graph. Edges compatible with the previously defined hub thresholds were designated as edge-derived hubs and subsequently classified as connector or provincial hubs based on their participation coefficients. To prioritize nodes that play the strongest integrative roles, candidate hubs were ranked according to a hub-size metric, defined as the weighted strength of each hubs’s connections to other high-centrality hubs. This measure captures not only how strongly a hub communicates overall, but specifically how strongly it interacts with other central hubs in the network.

##### 2.6.4.6. Community overlap

To quantify the extent to which nodes participate across multiple edge-defined communities, for each community we computed an entropy index following Faskowitz et al. (2020). For each node i, the distribution of its incident edges across the k identified communities was represented by p_ic_, the proportion of node i’s edges assigned to community c. The Shannon entropy of this distribution 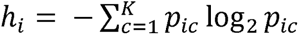 provides a measure of the diversity of community affiliations associated with node i. To enable comparison across communities, entropy values were normalized by their theoretical maximum, log_z_k, yielding a bounded measure:

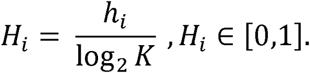

Higher values of H_i_ indicate that node’s edges are distributed more uniformly across communities, reflecting greater functional overlap and integrative connectivity, whereas lower values denote concentration within a single community, indicating more specialized or modular connectivity.

##### 2.6.4.7. Comparison between node-based and edge-based representations

To enable a direct comparison between time-averaged FC and time-varying FC, we projected the time-averaged FC matrices into edge space, following standard practice in edge-centric analyses (Faskowitz et al., 2020; Esfahlani et al., 2022). For each subject and run, upper-triangular elements of the time-averaged FC matrix were extracted to form edge-element vector representing all unique edges. These vectors were concatenated to compute an edge x edge covariance matrix by correlating each pair of edges across subjects and runs. This yielded a time-averaged edge-covariance matrix, directly comparable to the time-varying eFC matrix. Similarity between time-averaged and time-varying representations was quantified by correlating the upper triangles of both time-averaged and time-varying covariance matrices.

##### 2.6.4.8. Comparison between time-averaged and time-varying community structures

To quantify the similarity between different community partitions, we computed the Normalized Mutual Information (NMI) between: (i) the data-driven time-averaged community structure and the canonical Yeo-7 RSN labels, (ii) the time-varying community structure and RSN labels, and (iii) time-averaged and time-varying community partitions directly. NMI was computed using normalized_mutual_info_score from *scikit-learn* with arithmetic averaging, following the conventions of Alexander-Bloch et al. (2011), which corresponds to the Brain Connectivity Toolbox implementation. To assess statistical significance, we implemented a node-label permutation test. For each comparison, one partition was held fixed while the labels of the second partition were randomly permuted across nodes (preserving class-size distributions). We generated 5,000 permutations to build a null distribution of NMI values and computed one-sided p-values as the fraction of null NMIs ≥ the observed NMI. A standardized effect size (z-score) was computed as:

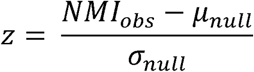

Null-distribution histograms were generated for visualization and quality control.

## 3. Results

We first validated our processing pipeline by replicating previously reported gastric-brain synchrony findings (Levakov et al., 2023a). To do so, Dice Similarity coefficient was calculated between our map and spatially permuted maps obtained from 10,000 spatial rotations of the Levakov et al. (2023a) map relative to our map. This resulted in a Dice similarity score = 0.536 *p* < .001, thus attesting to the validity of our methodology. Our analysis confirmed significant phase-locking between EGG and widely distributed cortical regions, consistent with established gastric network composition (Rebollo and Tallon-Baudry, 2022). In line with (Rebollo et al., 2018; Rebollo and Tallon-Baudry, 2022), we quantified the distribution of gastric network nodes across the Yeo-7 networks (Yeo et al., 2011; Figure 1A) and employed effect size (Cohen’ d) to statistically evaluate its anatomical composition. Forty regions emerged with moderate-high effect size (Figure 1B) including parcels across visual (striate and extrastriate), somatomotor, temporo-parietal, parietal (inferior and superior parietal lobules, intraparietal sulcus), prefrontal (medial/dorsal/ventral PFC), posterior cingulate, frontal operculum/insula, and occipito-temporal cortex (see Supplementary Table 1 for the full node list). nodes), visual (approximately 23%), salience, somatomotor, control, and dorsal attention networks (Rebollo and Tallon-Baudry, 2022). Additional hemispheric differences in gastric-network effect sizes and node distributions across Yeo-17 subnetworks are shown in Supplementary Figure S1. While this initial stage of network characterization revealed that the gastric network engages integration across resting-state networks, this characterization merely outlines the set of regions involved and their functional affiliations but does not yet inform us about the network’s organization or how it evolves over time.

**Figure 1.**
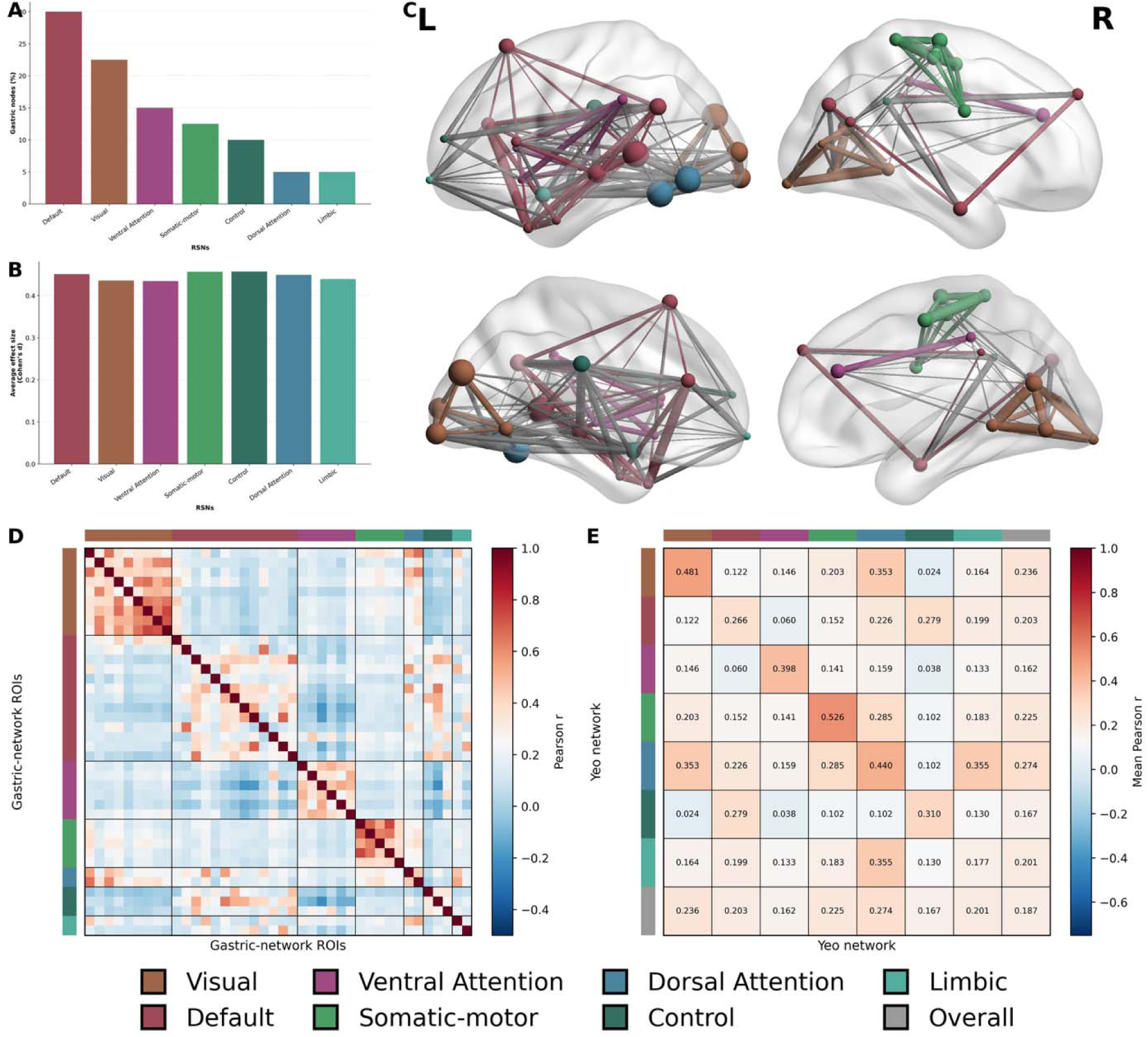
Anatomical and network composition of the time-averaged gastric network. (A) Distribution of gastric network nodes across the Yeo-7 networks, revealing a predominance of visual, ventral attention, and default mode networks. (B) Average effect size (Cohen’s d) of gastric nodes within each Yeo-7 network, showing comparable effect magnitudes across networks (Mean(SD) effect size across yeo-7 = .4461 (0.0096)). (C) Overlay of significant gastric–BOLD connections projected on the cortical surface of the right and left hemispheres shown in a lateral and medial view. Colors denote Yeo-7 resting-state network affiliation of the connected gastric nodes. Node size reflects degree centrality based on supra-threshold connections (|r| ≥ 0.20), scaled between 20–80 units (degree range 5–24). Edges represent correlations with |r| ≥ 0.20; line width increases with |r| strength, divided into four quantile bins (0.20–0.24, 0.24–0.31, 0.31–0.41, 0.41–0.75). (D) Functional connectivity matrix of gastric nodes ordered by their Yeo-7 resting-state network affiliation. (E) RSN-by-RSN average connectivity matrix showing the mean within- and between-network connectivity strengths. Patterns reflect strong intra-network interactions in sensory and default-mode regions. Heat scale shows group-level Pearson correlation coefficients (r) between gastric-network parcels, obtained by averaging Fisher z–transformed correlations across runs/subjects and back-transforming to r (blue = negative, red = positive).

### 3.1. Time-averaged Functional Connectivity Analysis of the Gastric Network

Moving beyond merely identifying individual brain regions synchronized with gastric activity, this analysis aimed to characterize the functional organization of these regions as a cohesive network. Graph theory provides a powerful framework to understand how these identified gastric-synchronized regions interact and act together as a unified system, revealing its overall structure and modularity. This time-averaged analysis is node-centric in its nature, as it treats brain regions as the irreducible units of brain structure and function (Faskowitz et al., 2020). The correlation matrix (Figure 1D) demonstrated that gastric-synchronized regions exhibited stronger functional connectivity within their respective resting-state networks than between different networks, consistent with prior evidence that interoceptive signals are embedded within intrinsic functional architectures (Tallon-Baudry, 2023). Within-network correlations (Figure 1E) were particularly high for regions associated with the visual network (*M=0.481*), consistent with previous findings showing that visual areas are prominent in the gastric network (Rebollo and Tallon-Baudry, 2022). Somatomotor network nodes also displayed increased internal connectivity (*M* = 0.526), forming a localized cluster. The ventral and dorsal attention modules also exhibited high inter-connectivity measures (*M* = 0.398 ; *M* = 0.44, respectively).

#### 3.1.1. Community structure within the gastric network

To characterize the internal organization of the network and to understand how this organization reflects its functional architecture, we applied the Louvain algorithm (Blondel et al., 2008), a widely used community-detection method for partitioning functional brain networks into cohesive functional modules. This approach identifies the partition that maximizes modularity, such that the resulting community structure represents the most stable and robust organization of the network.

After obtaining the optimal partition with a modularity value of *Q =* 0.58, the community analysis yielded 5 node-communities, presented in Figure 2. Community 1 was composed predominantly of DMN regions, with small additional contribution from control network. Community 2 consisted almost exclusively of visual regions, exhibiting a strong sensory profile with only minor involvement of DMN network. Community 3 encompassed core DMN territories alongside nodes from the dorsal attention, visual and limbic network, potentially reflecting an integrative association between internally oriented and sensory processes. Community 4 was exclusively dominated by nodes associated with the ventral attention network. A similar unimodal profile was also observed in community 5, with the involvement of only somatic module nodes.

**Figure 2.**
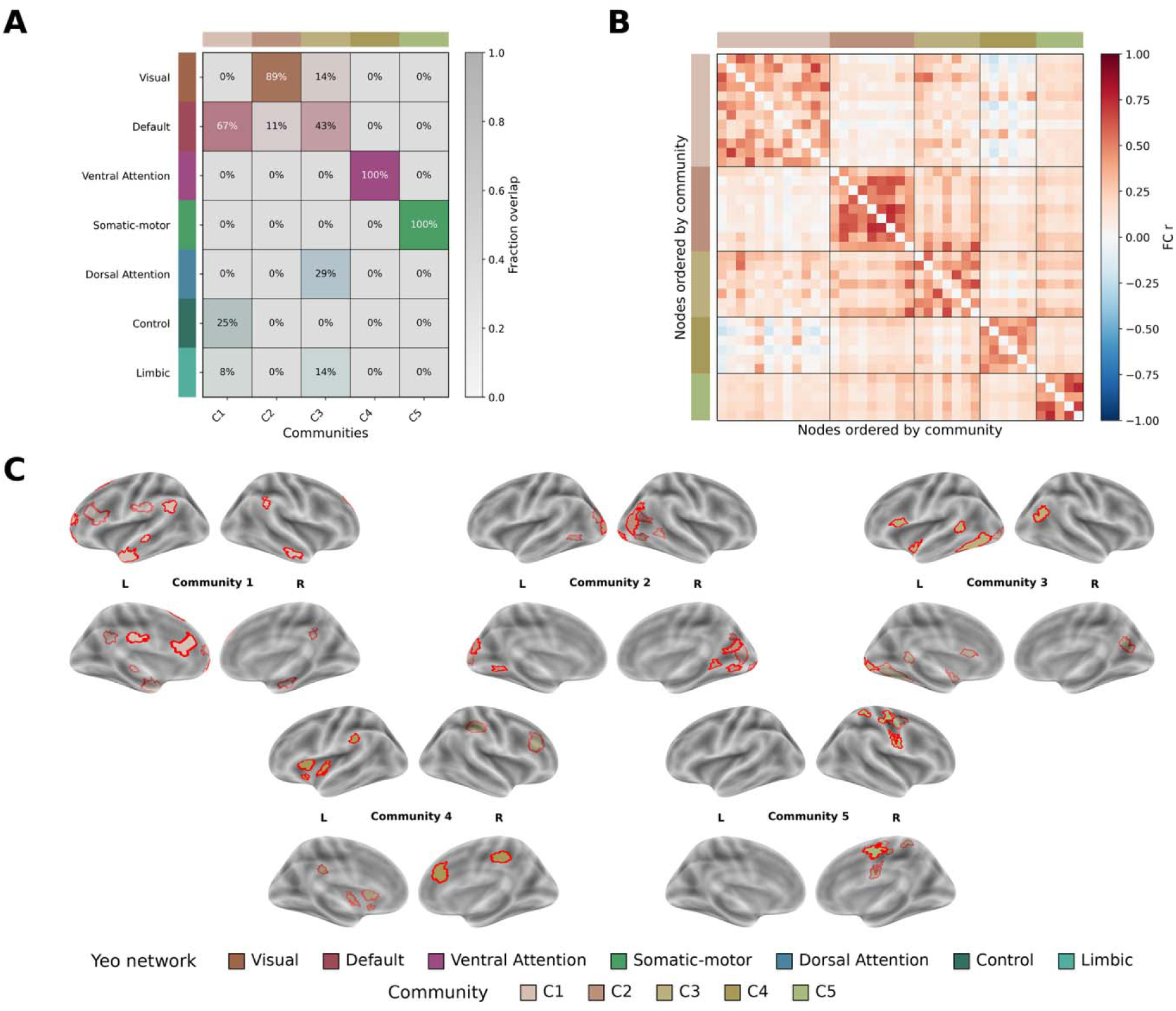
Spatial representation of gastric network communities and their correspondence with large-scale resting-state systems. (A) Fraction-overlap matrix quantifying the proportion of nodes shared between each community and each resting-state network (convention of color code of RSNs as in previous figures). Colored bars denote RSN membership, illustrating the modular organization underlying the gastric–BOLD network. Heat scale shows group-level Pearson correlation coefficients (r) between gastric-network communities, obtained by averaging Fisher z–transformed correlations across runs/subjects and back-transforming to r (blue = negative, red = positive). (B) The functional connectivity matrix of gastric nodes sorted by Louvain communities. (C) Surface maps shown in lateral and medial views show the cortical distribution of nodes belonging to each Louvain-derived community (Communities 1–5). Red contours denote the borders of gastric-synchronized regions within each community.

#### 3.1.2. Centrality measures

The time-averaged analysis detected ten hubs within the gastric network: four connector hubs and six provincial hubs (Figure 3). Connector hubs were defined as nodes of high weighted degree, alongside participation coefficients above 0.30, indicating strong involvement across multiple network communities. The strongest connector hub was located in the left extrastriate cortex within the visual network (node strength = 4.94, P = 0.49), followed by a left striate cortex hub (node strength = 4.81, P = 0.62). Two additional connector hubs were located in a left temporal subdivision of the default mode network (node strength = 4.10, P = 0.46) and in left temporo-occipital cortex within the dorsal attention network (node strength = 4.06, P = 0.63). In contrast, six provincial hubs were identified as high-strength nodes with p < 0.30 participation coefficients, suggesting a more locally embedded role within their respective communities. The strongest provincial hub was located in left inferior extrastriate cortex (node strength = 4.73, P = 0.26), followed by right inferior extrastriate cortex (node strength = 4.55, P = 0.00), left extrastriate cortex (node strength = 4.44, P = 0.15), right inferior extrastriate cortex (node strength = 4.44, P = 0.27), left superior extrastriate cortex (node strength = 4.35, P = 0.29), and a left inferior parietal region (node strength = 4.13, P = 0.00).

**Figure 3.**
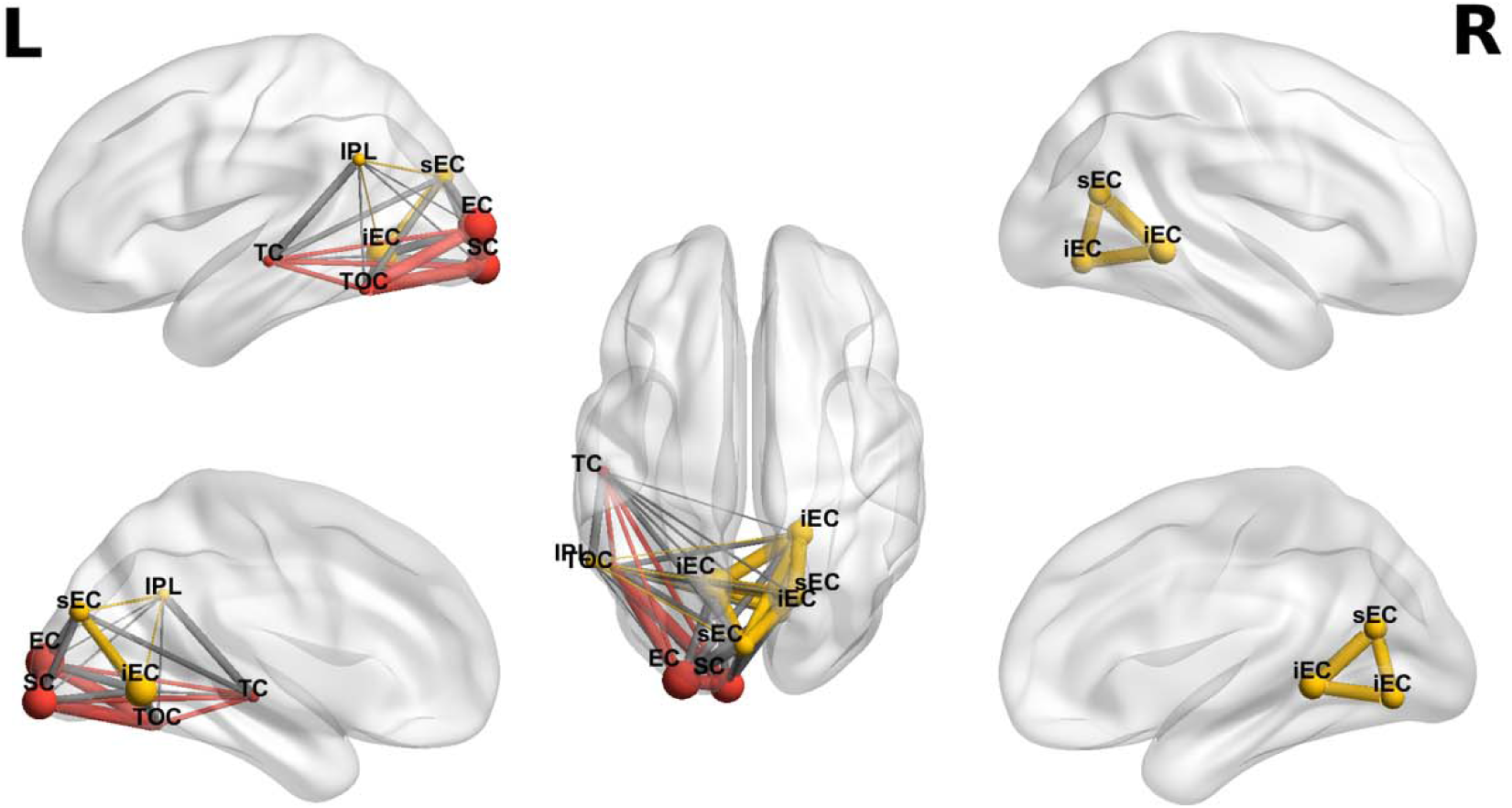
Time-averaged hub organization of the gastric network. Time-averaged hubs are displayed on inflated cortical surfaces shown from lateral, medial, and dorsal views. Connector hubs are shown in red and provincial hubs in yellow. Time-averaged connector hubs were restricted to the left hemisphere and were primarily centered on visual regions, including striate and extrastriate visual cortices, together with temporo-occipital cortex within the dorsal attention network and temporal cortex within the default mode network. Provincial hubs included peripheral visual regions in the superior and inferior extrastriate cortex, as well as inferior parietal lobule and a default-mode inferior parietal region. Abbreviations: IPL, inferior parietal lobule; sEC, superior extrastriate cortex; iEC, inferior extrastriate cortex; EC, extrastriate cortex; SC, striate cortex; TC, temporal cortex; TOC, temporo-occipital cortex; L, left hemisphere; R, right hemisphere.

### 3.2. Time-varying Functional Connectivity of the Gastric Network

Analysis of edge cofluctuations revealed time-varying, subject-specific burst patterns within the gastric network, indicating transient gastric-brain interactions (Betzel et al., 2023; Rebollo et al., 2018; Azzalini et al., 2019; see Supplementary Figure S2A). The combined empirical top-percentile and circular-shift thresholding procedure used to define burst frames is illustrated in Supplementary Figure S2B. As expected from eFC (Betzel et al., 2023), the overall structure and connectivity patterns from the time-varying analysis qualitatively resembled the time-averaged findings (Figure 4 and Figure 1 respectively).

**Figure 4.**
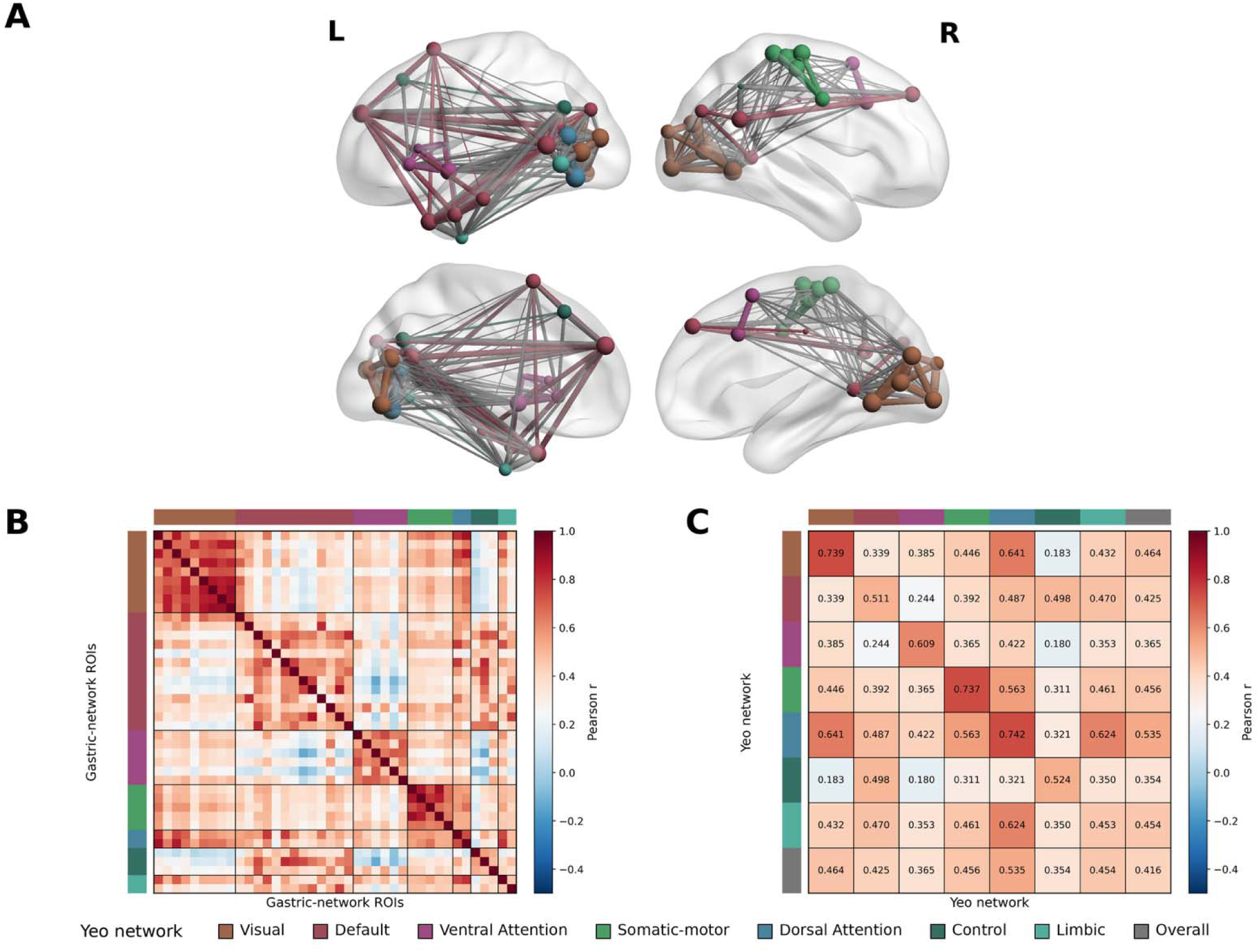
Dynamic, time-varying analysis of the large-scale functional architecture of the gastric network. (A) Spatial projection of the gastric-synchronized nodes and their connections on inflated cortical surfaces shown from lateral and medial views, color-coded by RSN affiliation. Node size reflects edge strength (sum of incident dynamic edge magnitudes), scaled from 10–100 radius units after robust clipping to the 5–95th percentile. Edge thickness reflects the magnitude of event-only edge-edge correlation above |r|>0.75 (B) RSN-by-RSN average connectivity matrix summarizing mean connectivity strengths, highlighting strong intra-network interactions and selective cross-network links among visual, somatomotor, ventral attention, and default networks. (C) Functional connectivity matrix of gastric edges ordered by Yeo-7 resting-state networks. Heat scale shows event-only edge correlation values (Fisher-z averaged across runs, then tanh-transformed back to r), ranging from −0.5 to 1.

To evaluate whether time-averaged and time-varying connectivity share a similar network structure, we correlated the upper triangles of the time-averaged edge covariance matrix and the time-varying EFC matrix using Pearson’s correlation. The similarity was modest but positive (*r* = 0.12, p < 0.0001), suggesting limited overlap between time-averaged and time-varying edge organization. Consistent with the time-averaged analysis, within-network correlations were especially pronounced in the somatomotor (*M* = 0.737) and visual (*M =* 0.739) networks, with dorsal attention nodes also displaying elevated internal connectivity (*M* = 0.742). The ventral attention network demonstrated robust interconnectivity (*M* = 0.609).

#### 3.2.1. Edge community structure and time-varying organization

Clustering the edge-centric functional connectivity (eFC) matrices revealed 6 time-varying communities encompassing the regions that constitute the gastric network (Figure 5). Each edge, representing the moment-to-moment co-fluctuation between a pair of brain regions, was assigned to a community and subsequently projected back onto its constituent nodes, permitting each region to participate in multiple community affiliations across its edges. The six identified communities exhibited distinct resting-state network (RSN) fingerprints, reflecting unique combinations of visual, attentional, somatomotor, limbic, and default-mode elements rather than direct replications of canonical RSNs. This observation suggests that the gastric network does not conform to a modular subdivision of established RSNs but instead comprises an integrated ensemble of functionally interactive systems. To further assess the robustness of the time-varying community structure to threshold selection, we quantified the similarity between community partitions obtained using the primary 5% burst threshold and those derived from more liberal thresholds (10%, 15%, and 25%) using normalized mutual information (NMI). Community structure was highly consistent across thresholds (5% vs. 10%: NMI = 0.792, p = 0.0002, z = 15.50; 5% vs. 15%: NMI = 0.792, p = 0.0002, z = 15.49; 5% vs. 25%: NMI = 0.792, p = 0.0002, z = 15.31), indicating that the identified time-varying organization is stable and not dependent on the specific threshold used for burst detection (see Supplementary Figure S3).

**Figure 5.**
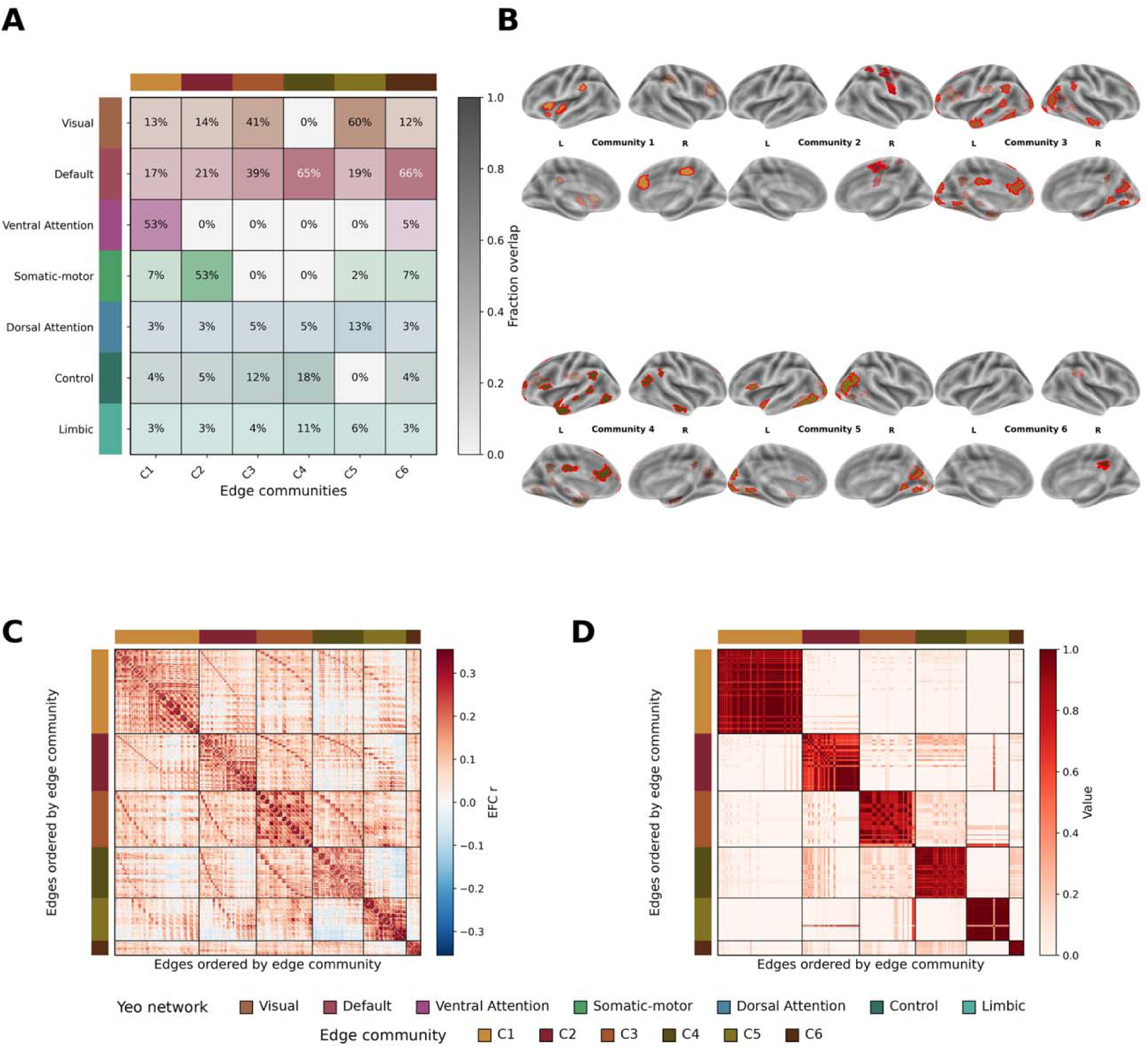
Spatial distribution of dynamic communities and their overlap with time-averaged modules. (A) Fraction-overlap matrix quantifying correspondence between edge-communities and RSN modules (convention of color code of RSNs as in previous figures). Colors on the matrix denote RSN membership. Color legends for edge-communities and yeo-7 RSN at the bottom of the figure. (B) Surface maps shown in a lateral and medial view show the cortical locations of each edge-community (Communities 1-6), with red highlights parcels involved in these edge-communities. (C) Edge-centric connectivity matrix sorted by edge-communities. Heat scale represents group-level event-only edge–edge correlations (*r*), obtained by averaging Fisher-*z* values across runs and back-transforming with *tanh*. Warmer colors indicate stronger positive connectivity; cooler colors indicate negative connectivity. (D) co-assignment probability matrix over 100 iterations of the Louvain algorithm. Colored bars indicate edge-community membership. Heat scale represents the probability that each edge will be assigned to each community in each iteration. Darker cells indicate edge pairs that are consistently grouped together and thus belong to more stable communities.

The edge-centric time-varying connectivity matrix (Figure 5C) demonstrated modularity value of Q = 0.44 and unfolded the networks into 6 edge-communities. Community 1 was composed primarily of ventral attention regions, followed by mainly DMN and visual contributions. Community 2 was dominated by somatomotor, with visual and DMN regions contributions. Community 3 showed combined dominance of visual and DMN sites. Community 4 exhibited a strong DMN dominant profile, with some contribution of the control network. Community 5 was characterized by visual regions as its principal constituents, accompanied by some contribution of the dorsal attention network. Finally, Community 6 showed a dominant DMN presence, followed by minimal involvement of visual and somatomotor regions. A key pattern observed in these overlapping communities is the substantial involvement of DMN and visual regions throughout multiple communities, in agreement with their prevalent presence in the time-varying gastric network. The phase-randomization control analysis showed that observed eFC was largely uncorrelated with surrogate eFC, supporting the interpretation that the observed time-varying structure is not explained by intrinsic temporal autocorrelation alone (Supplementary Figure S4).

#### 3.2.2. Centrality measures

By restricting the analysis to these burst-moments of heightened cofluctuation, we aimed to identify the nodes that are most central precisely during the periods that drive the network’s time-varying organization. The time-varying analysis identified 124 connector hubs (see full hub list in Supplementary Table 2), indicating widespread cross-community integration during burst events. Ranking them by the summed strength of all edges these regions connect, reduced the set to the most integrative connector hubs (Figure 6). In contrast to the time-averaged analysis, where connector hubs were mainly sensory-oriented and located exclusively in the left hemisphere, the time-varying analysis revealed a markedly different profile. Connector hubs were located in both hemispheres and included regions in the somatomotor, temporal, parietal, prefrontal and visual cortices, alongside precuneus, cingulate cortex, orbitofrontal cortex, insula. In addition to these connector hubs, the time-varying analysis identified strong provincial hubs. The strongest provincial hub linked the temporo-occipital cortex with the insula, followed by a hub linking the left insula with the right somatomotor cortex (see detailed connector and provincial hubs list in Supplementary Table 3).

**Figure 6.**
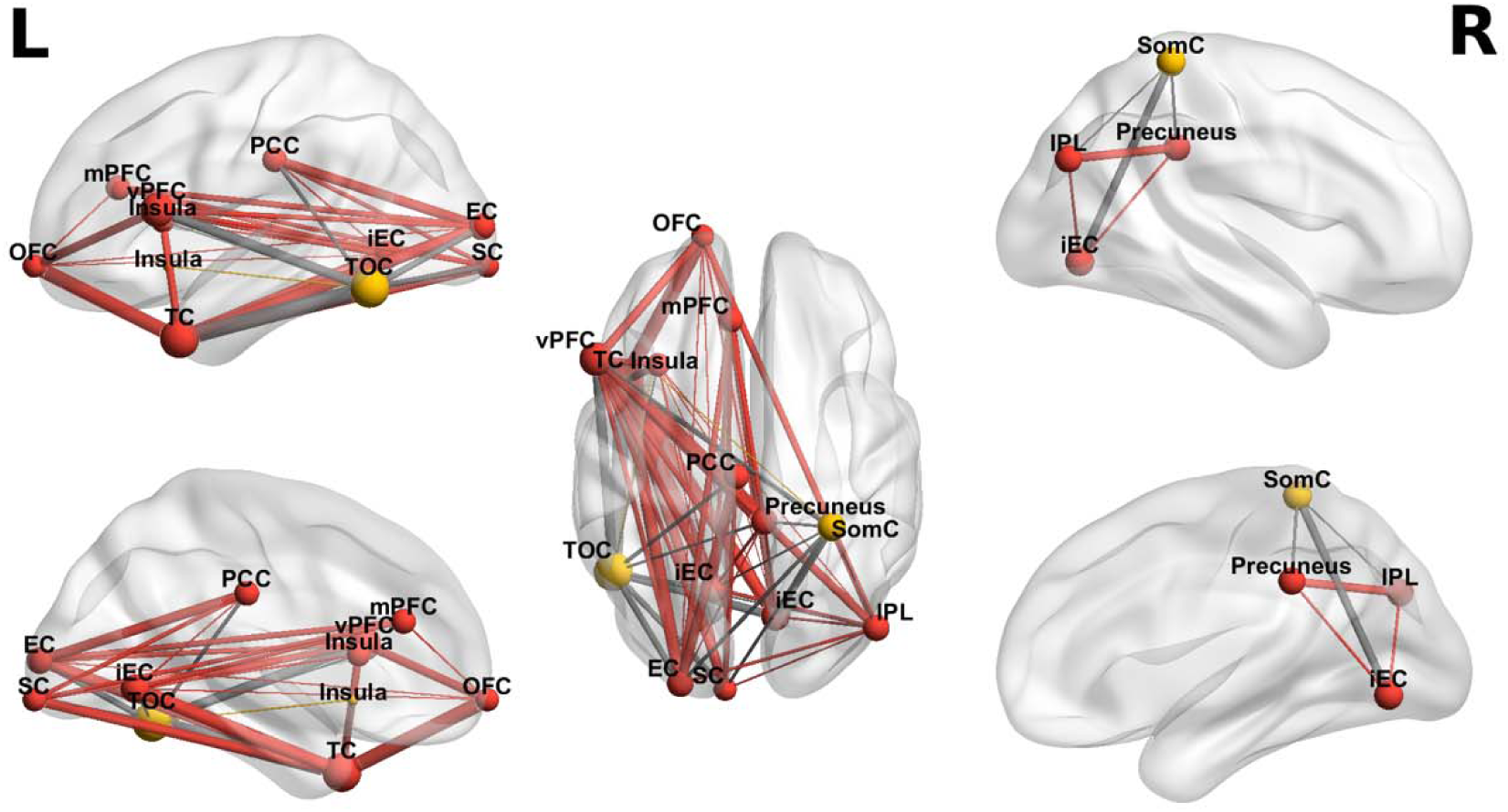
Edge-derived hub organization of the gastric network during burst events. Edge-derived hubs are displayed on inflated cortical surfaces from lateral, medial, and dorsal views. Connector hubs are shown in red and provincial hubs in yellow. In contrast to the time-averaged hub organization, connector hubs were distributed bilaterally and extended beyond visual cortices to include default-mode temporal cortex, ventral and medial prefrontal cortices, precuneus, inferior parietal lobule, dorsal-attention temporo-occipital cortex, control-network cingulate cortex, limbic orbitofrontal cortex, insula, and somatomotor cortex. Provincial hubs included temporo-occipital cortex, insula, and somatomotor cortex. L, left hemisphere; R, right hemisphere.

#### 3.2.3. Time-varying integration across RSNs

To further quantify the balance between specialization and integration within each community, we computed the normalized entropy of community assignments for every brain region, indexing the uniformity of its edge affiliations. Lower entropy values indicate a more modular and functionally specialized organization, whereas higher values denote greater overlap across systems. Entropy analyses demonstrated that communities 1 (mean = 0.587 ± 0.287) and 6 (mean = 0.618 ± 0.26) were characterized by lower entropy, consistent with a more modular architecture, whereas communities 2 (0.7 ± 0.16), 3 (0.76 ± 0.006), 4 (0.76 ± 0.003), and 5 (0.75 ± 0.07) showed higher entropy, reflecting a more distributed and integrative organizational profile.

#### 3.2.4. Time-averaged modularity vs. time-varying integration

To quantify the similarity between community structure and canonical resting-state networks (RSNs), we computed the normalized mutual information (NMI) between (i) time-averaged communities vs. RSN labels, (ii) time-varying communities vs. RSN labels, and (iii) time-averaged vs. time-varying communities. NMI was calculated using the arithmetic-mean variant of the normalized mutual information score (Alexander-Bloch et al., 2012), treating each node’s community/RSN assignment as a categorical label. For each comparison we report the observed NMI, a one-tailed p-value, and the standardized effect size. Node-communities showed a strong correspondence with RSN modular structure (*NMI* = 0.7047; *p* = .0002; *z* = 11.020), demonstrating that the time-averaged functional architecture is highly segregated and more closely aligned with established large-scale networks (Yeo et al., 2011). In contrast, edge-communities exhibited only modest correspondence with RSNs (*NMI* = 0.255; *p* = 0.104; *z* = 1.303), and this similarity did not exceed permutation-derived null expectations. This suggests that the time-varying coupling structure does not adhere to canonical RSN modularity, reflecting a community organization that is more integrated and cross-network in nature. Finally, node vs. edge communities showed very low mutual information (*NMI* = 0.1595; *p* = 0.5807; *z* = -0.301), indicating that the two approaches capture distinct network states. Null-distribution histograms for these community-structure comparisons are shown in Supplementary Figure S5.

## 4. Discussion

Previous studies delineated the components of the gastric network (Rebollo and Tallon-Baudry, 2022; Levakov, Ganor, et al., 2023), however the topological organization of these regions has remained largely unexplored. The current study examined the organizational patterns of the gastric network through both time-averaged and time-varying perspectives. Using a time-averaged analysis, we characterized the network’s common topology. Using a time-varying analysis, we investigated how its functional architecture reconfigures over time. By employing edge-centric functional connectivity (eFC) analysis and comparing it to traditional node-centric approach, we demonstrate that the gastric network dynamically alternates between modular and integrative states. This reconfiguration may support the coordination and alignment of its distinct functional components.

Interoceptive networks are organized according to principles of modularity and integration, while shifting through transient “events” of cofluctuations (Leão et al., 2025; Schaefer et al., 2014; Tsakiris and Critchley, 2016). This pattern is in-line with gastric-brain coupling, which expresses both a stable topological structure and transient reconfigurations (Richter et al., 2017). First, the gastric network is not a fixed anatomical entity but a time-varying functional system, whose organization unfolds dynamically across the gastric cycle (Rebollo et al., 2018). Rather than time-averaged connectivity, it follows a temporal sequence of activations aligned to the gastric rhythm (Rebollo and Tallon-Baudry, 2022). Second, rs-FC is increasingly understood as arising from temporally structured fluctuations in regional activity and inter-regional cofluctuation events, rather than from stationary time-averaged coupling alone (Bertolero et al., 2015). Recent work suggests that both local BOLD signal variability and global variability in tv-FC constitute biologically meaningful features of multiscale brain organization, rather than merely reflecting noise or scanner-related artifact (Baracchini et al., 2026). In line with this view, gastric-BOLD coupling shows slow temporal fluctuations that co-occur with changes in BOLD amplitude and emerge synchronously across gastric-network nodes (Rebollo et al., 2018). Thus, a comprehensive characterization of the gastric network necessitates considering both its time-averaged structure and its time-varying reorganization, while evaluating how both prisms reflect the interplay between segregation and large-scale integration that shapes its functional architecture.

### 4.1. Overlapping communities as a marker of integration

The time-averaged analysis revealed a stable scaffold, reflected in well-defined RSN-based modularity (Figure 1D). Strong within-network correlations were found in somatomotor and visual sub-networks (Figure 1E). The community structure revealed a segregated state of the network, with communities comprising a unimodal function. Only community 3 diverged, displaying a more heterogeneous composition of functional modalities (Figure 2). Interestingly, this organization suggests that the observed architecture is not simply a reflection of canonical RSN hierarchy but may reflect a flexible coordination of functional modules according to the network demands, as evidenced by the coexistence of multiple RSNs within specific communities. In contrast to the time-averaged analysis, time-varying edge-communities showed relatively little correspondence with predefined RSNs, indicating that reconfigurations in the network reflects a more integrated, cross-network state, unconstrained by time-averaged boundaries (Sporns, 2013). These findings were supported by the NMI similarity analysis which revealed a sharp dissociation between time-averaged and time-varying community structures (see Supplementary Figure S5).

### 4.2. Hubs’ detection reveals reconfiguration in network core

The time-averaged analysis revealed a hub organization comprising four connector hubs and six provincial hubs spanning sensory and associative territories. The connector hubs included striate and extrastriate cortices, temporo-occipital cortex, and temporal cortex, all localized to the left hemisphere (Figure 3). The prominence of occipital connector hubs is consistent with prior evidence showing gastric-BOLD phase synchronization across visual cortex and the positioning of the gastric network within low-level sensory gradients (Rebollo et al., 2018; Rebollo and Tallon-Baudry, 2022). Beyond sensory regions, the temporo-occipital connector hub links the gastric network to systems supporting perceptual selection and stimulus relevance; notably, it lies near recently identified food-selective visual regions in ventral temporal cortex, potentially situating it at the interface of visual perception and metabolic meaning (Bannert and Bartels, 2022; Jain et al., 2023; Khosla et al., 2022; Pennock et al., 2026). The temporal connector hub extends this pattern into multisensory and associative domains, consistent with its role in integration, interoceptive awareness, and broader gut-brain connectivity (Thye et al., 2017; Haruki and Ogawa, 2023; Mulder et al., 2023). In parallel, the six provincial hubs were primarily located in visual cortices, including bilateral inferior extrastriate, left extrastriate, left inferior parietal and inferior parietal cortices. This provincial pattern suggests that the time-averaged gastric network may relate to specialized sensory processing. Our time-varying analysis revealed an expansion of the network’s capacity, with 124 edges transiently adopting connector hub status, thus exhibiting a sharp divergence from the sparse connector configuration observed in the time-averaged connectome. This expansion in the number of connector hubs suggests that information exchange across gastric-related modules is not fixed, but rather emerges through a highly flexible, time-resolved topology (Bertolero et al., 2015). To distil the essential architecture of this system, we identified a stable “integrative core” of these connector hubs. Connector hubs included the temporal cortex, temporo-occipital cortex, ventral prefrontal cortex, bilateral inferior extrastriate cortices, precuneus, extrastriate cortex, inferior parietal lobule, somatomotor cortex, cingulate cortex, medial prefrontal cortex, orbitofrontal cortex, striate cortex, insula, and superior extrastriate cortex. The presence of the insula among the time-varying connector hubs is particularly notable, given its role as a primary viscerosensory relay region. As an interoceptive hub, the insula may broadcast gastric physiological signals to association cortices, supporting the integration of bodily states with motivational and action-oriented processes (Tallon-Baudry et al., 2026; Avery et al., 2015). Consistent with this interpretation, the involvement of the IPL, precuneus, medial and ventral prefrontal regions, and temporal DMN regions points to a distributed set of convergence zones in which exteroceptive and interoceptive information may be coordinated and integrated. The hub status of the right IPL aligns with its role in bodily self-awareness (Babo-Rebelo and Tallon-Baudry, 2018), whereas temporal and prefrontal DMN hubs may contribute to assigning emotional, contextual, and regulatory significance to visceral signals (Pehrs et al., 2015; Stern et al., 2017).

In addition to this connector architecture, the time-varying analysis also revealed a provincial hub-organization. The strongest provincial edge linked left temporo-occipital cortex with the left insula, while the second provincial hub edge connected the left insula with the right somatomotor cortex. Thus, while the connector hubs reflect broad cross-community integration during burst-events, the provincial edges suggest that these events also preserve strongly embedded relationships between attentional, interoceptive, and somatomotor components of the gastric network.

The dual-mode architecture of the gastric network we observed, aligns with hierarchical predictive coding models, which posit that the brain maintains allostatic regulation by integrating ascending visceral physiological signals with top-down contextual priors (Craig, 2003; Critchley and Harrison, 2013). In this framework, the time-averaged, unimodal phase may represent an anticipatory state, maintaining a baseline and early predictive model of the gut’s status through sensory anchored hubs. Conversely, the emergent edge-derived connector hubs may represent the “interpretation” phase, where sensory cues are fused with exteroceptive context to coordinate autonomic, endocrine, and metabolic responses (Schulkin and Sterling, 2019; Savoca et al., 2024). Our results provide a topological basis for this allostatic control. The dual mode we observed, indicated by rhythmically switching between a modular, sensory-focused state and a high-order integrative core, may be indicative of the brain’s capacity to integrate physiologial and sensory sampling to a unified interpretation. Specifically, the recruitment of multimodal regions like the IPL, insula, somatomotor cortex, precuneus, and DMN regions during integrative bursts, suggests a mechanism for contextualizing gastric signals within a higher and holistic interoceptive state (Azzalini et al., 2019). By alternating between these modes, the gastric network optimizes the trade-off between sensory precision, varying physiological states, and the complex coordination between them, required for interoceptive and allostatic regulation (Rebollo and Tallon-Baudry, 2022).

### 4.3. Limitations and future directions

While this study identified reconfigurations in the gastric network, related to sensory and integrative roles, resting-state data do not allow inference about the directionality of gastric-cortical interactions. Gastric cues could drive cortical reconfigurations, cortical states could shape gastric activity, or both could reflect modulation by a shared process (Wolpert et al., 2020; Lurie et al., 2019). To understand whether these reconfigurations reflect top-down, bottom-up influences, or reciprocal feedback, specific manipulations should be employed in the experimental design. Food/water intake or mechanical stimulation of the stomach could act as a “bottom-up” stimulus (Mayeli et al., 2023), whereas visual cues could reflect a “top-down” mechanism (Oseran et al., in press).

Additionally, the burst events have been detected in rest, thus could not be assigned with a functional meaning (Betzel et al., 2023). To assess if bursts were genuine dynamics, rather than statistical artifacts, a null model was generated by randomly shuffling each edge’s time series, preserving its distribution (Betzel et al., 2023; Faskowitz et al., 2021). However, this permutation assumes temporal independence, ignoring autocorrelation and spatial correlations between edges, posing a limitation by underestimating true burst dynamics. Recent evidence shows that low-frequency fMRI cofluctuations are tightly coupled with autonomic physiological signals across multiple bodily systems and may reflect arousal-related brain-body dynamics (Bolt et al., 2025). Thus, future work could address this limitation by incorporating synchronized physiological signals as temporal anchors.

## Data availability

The data supporting the findings of this study are openly available via figshare at https://doi.org/10.6084/m9.figshare.21952910. Custom analysis code is available at https://github.com/yeela-z/gastric-brain-network-analysis.

## CrediT authorship contribution statement

**Y.Z.:** Conceptualization, Methodology, Software, Formal analysis, Investigation, Data curation, Writing – original draft, Writing – review & editing, Visualization.

**G.A.:** Conceptualization, Methodology, Investigation, Writing – original draft, Writing – review & editing, Supervision, Project administration, Funding acquisition.

## Acknowledgments

This work was supported by the Israel Science Foundation (ISF) [Grant No. 2274/21] awarded to G.A. The funder had no role in the study design, data collection, analysis, interpretation, or the decision to publish this work.

## Declaration of generative AI use

During the preparation of this work, the author used ChatGPT to enhance the language, assist with reference organization and academic-style formatting, and support literature summarization and sorting. After using this tool, the author reviewed and edited the content as needed and takes full responsibility for the content of the publication.

