## Supplementary Materials for "Time-averaged and Time-varying Structure of the Gastric Network Revealed Through fMRI-Electrogastrogram Synchronization"

**Supplementary Table 1. Nodes of the gastric network defined from effect size-based parcel selection.**

The table contains the cortical regions comprising the gastric network obtained from the group level analysis. Nodes were defined by intersecting significant gastric–BOLD synchrony voxels (empirical > chance-level PLV) with the Schaefer 400-parcel atlas (2 mm, 17-network resolution; Schaefer et al., 2018). For each parcel, we report the Schaefer label, parcel index within the atlas, hemisphere, large-scale Yeo network label, subnetwork label, parcel index in the original Schaefer ordering, and atlas ID. Only parcels exceeding the median Cohen’s d across all significant voxels within that parcel, were retained and are listed in the table.

| label | parcel_id | hemi | net_base | net_sub | idx | atlas_id |
| --- | --- | --- | --- | --- | --- | --- |
| 17Networks_RH_DefaultC_IPL_2 | 387 | RH | Default | DefaultC | 2 | 386 |
| 17Networks_LH_TempPar_4 | 199 | LH | TempPar | TempPar | 4 | 198 |
| 17Networks_RH_SomMotA_7 | 231 | RH | SomMot | SomMotA | 7 | 230 |
| 17Networks_LH_ContC_Cingp_2 | 149 | LH | Cont | ContC | 2 | 148 |
| 17Networks_RH_VisPeri_ExStrInf_3 | 216 | RH | Vis | VisPeri | 3 | 215 |
| 17Networks_RH_SomMotA_5 | 229 | RH | SomMot | SomMotA | 5 | 228 |
| 17Networks_RH_SomMotB_Cent_2 | 258 | RH | SomMot | SomMotB | 2 | 257 |
| 17Networks_LH_DefaultA_PFCm_6 | 167 | LH | Default | DefaultA | 6 | 166 |
| 17Networks_RH_SomMotA_6 | 230 | RH | SomMot | SomMotA | 6 | 229 |
| 17Networks_LH_DefaultB_Temp_6 | 173 | LH | Default | DefaultB | 6 | 172 |
| 17Networks_LH_DefaultB_IPL_2 | 175 | LH | Default | DefaultB | 2 | 174 |
| 17Networks_LH_DorsAttnA_TempOcc_4 | 64 | LH | DorsAttn | DorsAttnA | 4 | 63 |
| 17Networks_RH_DefaultC_IPL_1 | 386 | RH | Default | DefaultC | 1 | 385 |
| 17Networks_LH_DefaultB_PFCd_5 | 180 | LH | Default | DefaultB | 5 | 179 |
| 17Networks_LH_DefaultB_Temp_2 | 169 | LH | Default | DefaultB | 2 | 168 |
| 17Networks_LH_DorsAttnA_TempOcc_3 | 63 | LH | DorsAttn | DorsAttnA | 3 | 62 |
| 17Networks_RH_SalVentAttnB_PFCmp_1 | 312 | RH | SalVentAttn | SalVentAttnB | 1 | 311 |
| 17Networks_RH_DefaultA_pCunPCC_3 | 366 | RH | Default | DefaultA | 3 | 365 |
| 17Networks_LH_SalVentAttnA_Ins_2 | 91 | LH | SalVentAttn | SalVentAttnA | 2 | 90 |
| 17Networks_LH_DefaultB_PFCv_5 | 188 | LH | Default | DefaultB | 5 | 187 |
| 17Networks_LH_LimbicB_OFC_5 | 114 | LH | Limbic | LimbicB | 5 | 113 |
| 17Networks_RH_VisCent_ExStr_6 | 207 | RH | Vis | VisCent | 6 | 206 |
| 17Networks_LH_LimbicA_TempPole_7 | 121 | LH | Limbic | LimbicA | 7 | 120 |
| 17Networks_RH_VisCent_ExStr_9 | 211 | RH | Vis | VisCent | 9 | 210 |
| 17Networks_RH_ContB_IPL_1 | 339 | RH | Cont | ContB | 1 | 338 |
| 17Networks_RH_VisPeri_ExStrSup_1 | 221 | RH | Vis | VisPeri | 1 | 220 |
| 17Networks_LH_VisCent_Striate_1 | 8 | LH | Vis | VisCent | 1 | 7 |
| 17Networks_LH_ContB_PFClv_3 | 143 | LH | Cont | ContB | 3 | 142 |
| 17Networks_LH_VisPeri_ExStrInf_2 | 15 | LH | Vis | VisPeri | 2 | 14 |
| 17Networks_RH_DefaultA_Temp_1 | 359 | RH | Default | DefaultA | 1 | 358 |
| 17Networks_LH_SalVentAttnB_Ins_1 | 105 | LH | SalVentAttn | SalVentAttnB | 1 | 104 |
| 17Networks_RH_SomMotA_9 | 233 | RH | SomMot | SomMotA | 9 | 232 |
| 17Networks_LH_SalVentAttnA_Ins_3 | 92 | LH | SalVentAttn | SalVentAttnA | 3 | 91 |
| 17Networks_RH_VisPeri_ExStrInf_5 | 218 | RH | Vis | VisPeri | 5 | 217 |
| 17Networks_LH_VisPeri_ExStrSup_2 | 22 | LH | Vis | VisPeri | 2 | 21 |
| 17Networks_LH_SalVentAttnA_ParOper_3 | 89 | LH | SalVentAttn | SalVentAttnA | 3 | 88 |
| 17Networks_LH_VisCent_ExStr_8 | 10 | LH | Vis | VisCent | 8 | 9 |
| 17Networks_RH_SalVentAttnA_ParMed_2 | 299 | RH | SalVentAttn | SalVentAttnA | 2 | 298 |
| 17Networks_LH_DefaultB_Temp_1 | 168 | LH | Default | DefaultB | 1 | 167 |
| 17Networks_RH_DefaultB_PFCd_2 | 379 | RH | Default | DefaultB | 2 | 378 |

**Supplementary Table 2:** **Dynamic connector edge-derived hubs in the gastric network.**
The table presents all edge-hubs classified as connector hubs in the dynamic edge-centric analysis. Connector hubs were defined as edges that passed the strength-hub criterion and showed high participation coefficient across edge communities, $P>0.30$. Each row represents one edge hub, defined by the pair of Schaefer 17-network parcels forming that edge (node i, j). For each edge, the table reports its edge-graph degree, edge strength, and participation coefficient $P$. A total of 124 connector edge-derived hubs were identified, reflecting transiently integrative edges that linked multiple functional communities during high co-fluctuation states.

| Node i | Node j | Degree | Strength | P |
| --- | --- | --- | --- | --- |
| 17Networks_LH_ContC_Cingp_2 | 17Networks_LH_DorsAttnA_TempOcc_3 | 209 | 45.374 | 0.656 |
| 17Networks_LH_ContC_Cingp_2 | 17Networks_LH_DorsAttnA_TempOcc_4 | 158 | 36.551 | 0.630 |
| 17Networks_LH_ContC_Cingp_2 | 17Networks_LH_LimbicA_TempPole_7 | 169 | 37.075 | 0.656 |
| 17Networks_LH_ContC_Cingp_2 | 17Networks_LH_VisCent_Striate_1 | 180 | 39.455 | 0.663 |
| 17Networks_LH_ContC_Cingp_2 | 17Networks_LH_VisPeri_ExStrInf_2 | 151 | 35.842 | 0.352 |
| 17Networks_LH_ContC_Cingp_2 | 17Networks_LH_VisPeri_ExStrSup_2 | 179 | 39.539 | 0.612 |
| 17Networks_LH_ContC_Cingp_2 | 17Networks_LH_VisCent_ExStr_8 | 214 | 47.069 | 0.635 |
| 17Networks_LH_ContC_Cingp_2 | 17Networks_RH_VisPeri_ExStrSup_1 | 173 | 40.201 | 0.465 |
| 17Networks_LH_ContC_Cingp_2 | 17Networks_RH_VisPeri_ExStrInf_3 | 149 | 34.366 | 0.435 |
| 17Networks_LH_ContC_Cingp_2 | 17Networks_RH_VisPeri_ExStrInf_5 | 161 | 38.091 | 0.394 |
| 17Networks_LH_ContC_Cingp_2 | 17Networks_LH_DorsAttnA_TempOcc_3 | 180 | 38.693 | 0.722 |
| 17Networks_RH_ContB_IPL_1 | 17Networks_RH_DefaultA_pCunPCC_3 | 156 | 36.264 | 0.659 |
| 17Networks_RH_ContB_IPL_1 | 17Networks_LH_DorsAttnA_TempOcc_3 | 178 | 40.249 | 0.674 |
| 17Networks_RH_ContB_IPL_1 | 17Networks_LH_LimbicB_OFC_5 | 174 | 37.640 | 0.703 |
| 17Networks_RH_ContB_IPL_1 | 17Networks_LH_VisCent_Striate_1 | 157 | 36.280 | 0.378 |
| 17Networks_RH_ContB_IPL_1 | 17Networks_LH_VisPeri_ExStrInf_2 | 155 | 35.152 | 0.550 |
| 17Networks_RH_ContB_IPL_1 | 17Networks_LH_VisPeri_ExStrSup_2 | 170 | 38.176 | 0.557 |
| 17Networks_RH_ContB_IPL_1 | 17Networks_LH_VisCent_ExStr_8 | 155 | 36.878 | 0.437 |
| 17Networks_RH_ContB_IPL_1 | 17Networks_RH_VisPeri_ExStrSup_1 | 150 | 34.709 | 0.484 |
| 17Networks_RH_ContB_IPL_1 | 17Networks_RH_VisPeri_ExStrInf_3 | 145 | 35.009 | 0.351 |
| 17Networks_LH_DefaultB_Temp_1 | 17Networks_RH_DefaultA_pCunPCC_3 | 260 | 55.620 | 0.679 |
| 17Networks_LH_DefaultB_Temp_1 | 17Networks_LH_DorsAttnA_TempOcc_3 | 259 | 55.504 | 0.695 |
| 17Networks_LH_DefaultB_Temp_1 | 17Networks_LH_LimbicB_OFC_5 | 249 | 53.501 | 0.718 |
| 17Networks_LH_DefaultB_Temp_1 | 17Networks_LH_LimbicA_TempPole_7 | 184 | 40.709 | 0.567 |
| 17Networks_LH_DefaultB_Temp_1 | 17Networks_LH_SalVentAttnB_Ins_1 | 176 | 38.583 | 0.374 |
| 17Networks_LH_DefaultB_Temp_1 | 17Networks_LH_SalVentAttnA_Ins_3 | 198 | 43.846 | 0.563 |
| 17Networks_LH_DefaultB_Temp_1 | 17Networks_LH_SalVentAttnA_ParOper_3 | 203 | 45.390 | 0.633 |
| 17Networks_LH_DefaultB_Temp_1 | 17Networks_RH_SalVentAttnB_PFCmp_1 | 187 | 40.735 | 0.643 |
| 17Networks_LH_DefaultB_Temp_1 | 17Networks_RH_SomMotA_5 | 157 | 34.691 | 0.635 |
| 17Networks_LH_DefaultB_Temp_1 | 17Networks_RH_SomMotA_7 | 166 | 37.150 | 0.684 |
| 17Networks_LH_DefaultB_Temp_1 | 17Networks_RH_SomMotA_9 | 182 | 40.769 | 0.646 |
| 17Networks_LH_DefaultB_Temp_1 | 17Networks_LH_TempPar_4 | 237 | 51.311 | 0.670 |
| 17Networks_LH_DefaultB_Temp_1 | 17Networks_LH_VisCent_Striate_1 | 203 | 47.531 | 0.500 |
| 17Networks_LH_DefaultB_Temp_1 | 17Networks_LH_VisPeri_ExStrInf_2 | 239 | 52.553 | 0.637 |
| 17Networks_LH_DefaultB_Temp_1 | 17Networks_LH_VisPeri_ExStrSup_2 | 233 | 51.742 | 0.591 |
| 17Networks_LH_DefaultB_Temp_1 | 17Networks_LH_VisCent_ExStr_8 | 218 | 50.240 | 0.530 |
| 17Networks_LH_DefaultB_Temp_1 | 17Networks_RH_VisPeri_ExStrSup_1 | 217 | 48.631 | 0.557 |
| 17Networks_LH_DefaultB_Temp_1 | 17Networks_RH_VisPeri_ExStrInf_3 | 215 | 50.901 | 0.534 |
| 17Networks_LH_DefaultB_Temp_1 | 17Networks_RH_VisPeri_ExStrInf_5 | 207 | 47.183 | 0.537 |
| 17Networks_LH_DefaultB_Temp_2 | 17Networks_LH_DorsAttnA_TempOcc_3 | 168 | 35.007 | 0.742 |
| 17Networks_LH_DefaultB_PFCv_5 | 17Networks_RH_DefaultA_pCunPCC_3 | 187 | 42.589 | 0.687 |
| 17Networks_LH_DefaultB_PFCv_5 | 17Networks_LH_DorsAttnA_TempOcc_3 | 213 | 47.033 | 0.701 |
| 17Networks_LH_DefaultB_PFCv_5 | 17Networks_LH_LimbicB_OFC_5 | 211 | 45.500 | 0.740 |
| 17Networks_LH_DefaultB_PFCv_5 | 17Networks_LH_LimbicA_TempPole_7 | 173 | 37.083 | 0.565 |
| 17Networks_LH_DefaultB_PFCv_5 | 17Networks_LH_SalVentAttnA_Ins_3 | 190 | 41.639 | 0.621 |
| 17Networks_LH_DefaultB_PFCv_5 | 17Networks_LH_SalVentAttnA_ParOper_3 | 196 | 41.427 | 0.647 |
| 17Networks_LH_DefaultB_PFCv_5 | 17Networks_RH_SalVentAttnB_PFCmp_1 | 165 | 35.013 | 0.654 |
| 17Networks_LH_DefaultB_PFCv_5 | 17Networks_RH_SomMotA_9 | 161 | 35.612 | 0.667 |
| 17Networks_LH_DefaultB_PFCv_5 | 17Networks_LH_TempPar_4 | 155 | 35.717 | 0.677 |
| 17Networks_LH_DefaultB_PFCv_5 | 17Networks_LH_VisCent_Striate_1 | 167 | 38.955 | 0.448 |
| 17Networks_LH_DefaultB_PFCv_5 | 17Networks_LH_VisPeri_ExStrInf_2 | 191 | 42.458 | 0.617 |
| 17Networks_LH_DefaultB_PFCv_5 | 17Networks_LH_VisPeri_ExStrSup_2 | 184 | 40.773 | 0.588 |
| 17Networks_LH_DefaultB_PFCv_5 | 17Networks_LH_VisCent_ExStr_8 | 177 | 40.174 | 0.530 |
| 17Networks_LH_DefaultB_PFCv_5 | 17Networks_RH_VisPeri_ExStrSup_1 | 165 | 37.648 | 0.574 |
| 17Networks_LH_DefaultB_PFCv_5 | 17Networks_RH_VisPeri_ExStrInf_3 | 176 | 40.740 | 0.506 |
| 17Networks_LH_DefaultB_PFCv_5 | 17Networks_RH_VisPeri_ExStrInf_5 | 169 | 38.079 | 0.471 |
| 17Networks_LH_DefaultA_PFCm_6 | 17Networks_RH_DefaultA_pCunPCC_3 | 178 | 39.643 | 0.699 |
| 17Networks_LH_DefaultA_PFCm_6 | 17Networks_LH_LimbicB_OFC_5 | 160 | 35.884 | 0.736 |
| 17Networks_LH_DefaultA_PFCm_6 | 17Networks_LH_SalVentAttnA_ParOper_3 | 177 | 37.205 | 0.562 |
| 17Networks_LH_DefaultA_PFCm_6 | 17Networks_LH_TempPar_4 | 158 | 35.837 | 0.703 |
| 17Networks_LH_DefaultA_PFCm_6 | 17Networks_LH_VisCent_Striate_1 | 170 | 39.279 | 0.553 |
| 17Networks_LH_DefaultA_PFCm_6 | 17Networks_LH_VisPeri_ExStrInf_2 | 168 | 38.786 | 0.677 |
| 17Networks_LH_DefaultA_PFCm_6 | 17Networks_LH_VisPeri_ExStrSup_2 | 167 | 38.544 | 0.654 |
| 17Networks_LH_DefaultA_PFCm_6 | 17Networks_LH_VisCent_ExStr_8 | 179 | 41.690 | 0.591 |
| 17Networks_LH_DefaultA_PFCm_6 | 17Networks_RH_VisPeri_ExStrSup_1 | 162 | 37.261 | 0.610 |
| 17Networks_LH_DefaultA_PFCm_6 | 17Networks_RH_VisPeri_ExStrInf_3 | 197 | 44.718 | 0.573 |
| 17Networks_LH_DefaultA_PFCm_6 | 17Networks_RH_VisPeri_ExStrInf_5 | 171 | 38.416 | 0.559 |
| 17Networks_RH_DefaultC_IPL_1 | 17Networks_RH_DefaultA_pCunPCC_3 | 201 | 43.269 | 0.637 |
| 17Networks_RH_DefaultC_IPL_1 | 17Networks_LH_LimbicB_OFC_5 | 191 | 40.819 | 0.713 |
| 17Networks_RH_DefaultC_IPL_1 | 17Networks_LH_VisCent_Striate_1 | 163 | 37.676 | 0.420 |
| 17Networks_RH_DefaultC_IPL_1 | 17Networks_LH_VisPeri_ExStrInf_2 | 171 | 37.454 | 0.575 |
| 17Networks_RH_DefaultC_IPL_1 | 17Networks_LH_VisPeri_ExStrSup_2 | 198 | 43.031 | 0.603 |
| 17Networks_RH_DefaultC_IPL_1 | 17Networks_LH_VisCent_ExStr_8 | 170 | 39.389 | 0.469 |
| 17Networks_RH_DefaultC_IPL_1 | 17Networks_RH_VisPeri_ExStrInf_3 | 166 | 38.950 | 0.424 |
| 17Networks_RH_DefaultC_IPL_1 | 17Networks_RH_VisPeri_ExStrInf_5 | 156 | 36.096 | 0.432 |
| 17Networks_RH_DefaultC_IPL_2 | 17Networks_RH_DefaultA_pCunPCC_3 | 203 | 44.793 | 0.663 |
| 17Networks_RH_DefaultC_IPL_2 | 17Networks_LH_DorsAttnA_TempOcc_3 | 159 | 35.791 | 0.637 |
| 17Networks_RH_DefaultC_IPL_2 | 17Networks_LH_LimbicB_OFC_5 | 198 | 42.517 | 0.718 |
| 17Networks_RH_DefaultC_IPL_2 | 17Networks_LH_SalVentAttnA_Ins_3 | 161 | 35.059 | 0.609 |
| 17Networks_RH_DefaultC_IPL_2 | 17Networks_RH_SomMotA_9 | 164 | 35.926 | 0.614 |
| 17Networks_RH_DefaultC_IPL_2 | 17Networks_LH_TempPar_4 | 173 | 38.630 | 0.657 |
| 17Networks_RH_DefaultC_IPL_2 | 17Networks_LH_VisCent_Striate_1 | 155 | 35.982 | 0.459 |
| 17Networks_RH_DefaultC_IPL_2 | 17Networks_LH_VisPeri_ExStrInf_2 | 174 | 38.608 | 0.633 |
| 17Networks_RH_DefaultC_IPL_2 | 17Networks_LH_VisPeri_ExStrSup_2 | 179 | 39.983 | 0.594 |
| 17Networks_RH_DefaultC_IPL_2 | 17Networks_LH_VisCent_ExStr_8 | 165 | 37.704 | 0.513 |
| 17Networks_RH_DefaultC_IPL_2 | 17Networks_RH_VisPeri_ExStrInf_3 | 157 | 37.006 | 0.459 |
| 17Networks_RH_DefaultC_IPL_2 | 17Networks_RH_VisPeri_ExStrInf_5 | 164 | 37.249 | 0.529 |
| 17Networks_RH_DefaultA_pCunPCC_3 | 17Networks_LH_DorsAttnA_TempOcc_3 | 160 | 36.893 | 0.721 |
| 17Networks_RH_DefaultA_pCunPCC_3 | 17Networks_LH_SalVentAttnA_Ins_3 | 158 | 34.942 | 0.440 |
| 17Networks_RH_DefaultA_pCunPCC_3 | 17Networks_RH_SomMotA_9 | 169 | 37.191 | 0.771 |
| 17Networks_RH_DefaultA_pCunPCC_3 | 17Networks_LH_VisPeri_ExStrInf_2 | 162 | 37.583 | 0.564 |
| 17Networks_RH_DefaultA_pCunPCC_3 | 17Networks_LH_VisCent_ExStr_8 | 144 | 35.575 | 0.488 |
| 17Networks_RH_DefaultA_pCunPCC_3 | 17Networks_RH_VisPeri_ExStrInf_3 | 146 | 36.970 | 0.465 |
| 17Networks_LH_DorsAttnA_TempOcc_3 | 17Networks_LH_LimbicA_TempPole_7 | 152 | 34.316 | 0.371 |
| 17Networks_LH_DorsAttnA_TempOcc_3 | 17Networks_LH_SalVentAttnA_Ins_3 | 156 | 35.636 | 0.399 |
| 17Networks_LH_DorsAttnA_TempOcc_3 | 17Networks_LH_SalVentAttnA_ParOper_3 | 177 | 37.921 | 0.388 |
| 17Networks_LH_DorsAttnA_TempOcc_3 | 17Networks_RH_SomMotA_7 | 157 | 34.869 | 0.531 |
| 17Networks_LH_DorsAttnA_TempOcc_3 | 17Networks_LH_VisCent_Striate_1 | 179 | 41.320 | 0.582 |
| 17Networks_LH_DorsAttnA_TempOcc_3 | 17Networks_LH_VisPeri_ExStrInf_2 | 161 | 37.135 | 0.680 |
| 17Networks_LH_DorsAttnA_TempOcc_3 | 17Networks_LH_VisCent_ExStr_8 | 182 | 42.028 | 0.622 |
| 17Networks_LH_DorsAttnA_TempOcc_3 | 17Networks_RH_VisPeri_ExStrSup_1 | 157 | 36.519 | 0.601 |
| 17Networks_LH_DorsAttnA_TempOcc_3 | 17Networks_RH_VisPeri_ExStrInf_3 | 194 | 45.198 | 0.607 |
| 17Networks_LH_DorsAttnA_TempOcc_3 | 17Networks_RH_VisPeri_ExStrInf_5 | 188 | 42.051 | 0.593 |
| 17Networks_LH_LimbicB_OFC_5 | 17Networks_LH_VisPeri_ExStrInf_2 | 156 | 35.153 | 0.646 |
| 17Networks_LH_LimbicB_OFC_5 | 17Networks_LH_VisCent_ExStr_8 | 157 | 35.301 | 0.554 |
| 17Networks_LH_LimbicB_OFC_5 | 17Networks_RH_VisPeri_ExStrSup_1 | 160 | 35.450 | 0.588 |
| 17Networks_LH_LimbicB_OFC_5 | 17Networks_RH_VisPeri_ExStrInf_3 | 159 | 36.488 | 0.538 |
| 17Networks_LH_SalVentAttnA_Ins_3 | 17Networks_RH_SomMotA_7 | 165 | 35.715 | 0.452 |
| 17Networks_LH_SalVentAttnA_Ins_3 | 17Networks_LH_VisCent_Striate_1 | 155 | 34.822 | 0.521 |
| 17Networks_LH_SalVentAttnA_Ins_3 | 17Networks_LH_VisPeri_ExStrInf_2 | 159 | 35.482 | 0.441 |
| 17Networks_LH_SalVentAttnA_Ins_3 | 17Networks_LH_VisPeri_ExStrSup_2 | 162 | 35.796 | 0.450 |
| 17Networks_LH_SalVentAttnA_Ins_3 | 17Networks_RH_VisPeri_ExStrInf_3 | 154 | 34.570 | 0.529 |
| 17Networks_LH_SalVentAttnA_ParOper_3 | 17Networks_RH_SomMotA_9 | 168 | 35.701 | 0.450 |
| 17Networks_RH_SomMotA_9 | 17Networks_LH_VisCent_Striate_1 | 173 | 37.613 | 0.667 |
| 17Networks_RH_SomMotA_9 | 17Networks_LH_VisPeri_ExStrInf_2 | 167 | 36.301 | 0.733 |
| 17Networks_RH_SomMotA_9 | 17Networks_LH_VisPeri_ExStrSup_2 | 185 | 39.430 | 0.744 |
| 17Networks_RH_SomMotA_9 | 17Networks_LH_VisCent_ExStr_8 | 176 | 38.591 | 0.691 |
| 17Networks_RH_SomMotA_9 | 17Networks_RH_VisPeri_ExStrSup_1 | 172 | 37.109 | 0.710 |
| 17Networks_RH_SomMotA_9 | 17Networks_RH_VisPeri_ExStrInf_3 | 203 | 44.382 | 0.680 |
| 17Networks_RH_SomMotA_9 | 17Networks_RH_VisPeri_ExStrInf_5 | 175 | 37.690 | 0.668 |
| 17Networks_LH_TempPar_4 | 17Networks_LH_VisPeri_ExStrInf_2 | 160 | 36.624 | 0.608 |
| 17Networks_LH_TempPar_4 | 17Networks_LH_VisCent_ExStr_8 | 144 | 34.926 | 0.536 |
| 17Networks_LH_TempPar_4 | 17Networks_RH_VisPeri_ExStrSup_1 | 146 | 34.365 | 0.537 |
| 17Networks_LH_TempPar_4 | 17Networks_RH_VisPeri_ExStrInf_3 | 162 | 40.249 | 0.549 |

**Supplementary Table 3: Top edge-derived hubs in the gastric network.**

This table presents both the connector and provincial hubs of the time-varying analysis. The core connector hubs are presented out of total of 124. Top connector hubs are “hubs of hubs” – hubs that are most connected to other connector hubs in the network. This procedure yielded 15 core connector hubs and 3 provincial hubs, detailed in the table below. Note that for clearer visualization and finer resolution of network distribution, here we present the yeo-17 RSN network subdivisions, and not the canonical yeo-7 used throughout the paper. Network labels follow the Yeo 17-network cortical parcellation: VisCent and VisPeri denote central and peripheral visual networks; SomMotA-B, somatomotor networks; DorsAttnA-B, dorsal attention networks; SalVentAttnA-B, salience/ventral attention networks; ContA-C, control networks; DefaultA-C, default mode networks; and LimbicA-B, limbic networks.

| Hub subdivision | node_idx | Label | Degree | Strength sum | Combined rank |
| --- | --- | --- | --- | --- | --- |
| **Provincial** | 19 | 17Networks_LH_SalVentAttnB_Ins_1 | 2.0 | 70.84 | 2 |
| **Provincial** | 15 | 17Networks_LH_DorsAttnA_TempOcc_3 | 1.0 | 35.60 | 4 |
| **Provincial** | 29 | 17Networks_RH_SomMotA_9 | 1.0 | 35.23 | 5 |
| **Connector** | 3 | 17Networks_LH_DefaultB_Temp_1 | 19.0 | 886.59 | 2 |
| **Connector** | 15 | 17Networks_LH_DorsAttnA_TempOcc_3 | 18.0 | 721.54 | 4 |
| **Connector** | 7 | 17Networks_LH_DefaultB_PFCv_5 | 16.0 | 640.44 | 6 |
| **Connector** | 36 | 17Networks_RH_VisPeri_ExStrInf_3 | 13.0 | 519.55 | 8 |
| **Connector** | 32 | 17Networks_LH_VisPeri_ExStrInf_2 | 13.0 | 499.13 | 9 |
| **Connector** | 14 | 17Networks_RH_DefaultA_pCunPCC_3 | 12.0 | 481.33 | 11 |
| **Connector** | 34 | 17Networks_LH_VisCent_ExStr_8 | 12.0 | 479.57 | 12 |
| **Connector** | 12 | 17Networks_RH_DefaultC_IPL_2 | 12.0 | 459.25 | 13 |
| **Connector** | 29 | 17Networks_RH_SomMotA_9 | 12.0 | 456.31 | 14 |
| **Connector** | 0 | 17Networks_LH_ContC_Cingp_2 | 11.0 | 432.26 | 16 |
| **Connector** | 8 | 17Networks_LH_DefaultA_PFCm_6 | 11.0 | 427.26 | 17 |
| **Connector** | 17 | 17Networks_LH_LimbicB_OFC_5 | 10.0 | 398.25 | 19 |
| **Connector** | 31 | 17Networks_LH_VisCent_Striate_1 | 10.0 | 388.91 | 20 |
| **Connector** | 21 | 17Networks_LH_SalVentAttnA_Ins_3 | 10.0 | 367.51 | 21 |
| **Connector** | 33 | 17Networks_LH_VisPeri_ExStrSup_2 | 9.0 | 367.01 | 23 |

**Supplementary Figure S1:** **Hemispheric differences in gastric-network properties across large-scale cortical subnetworks**

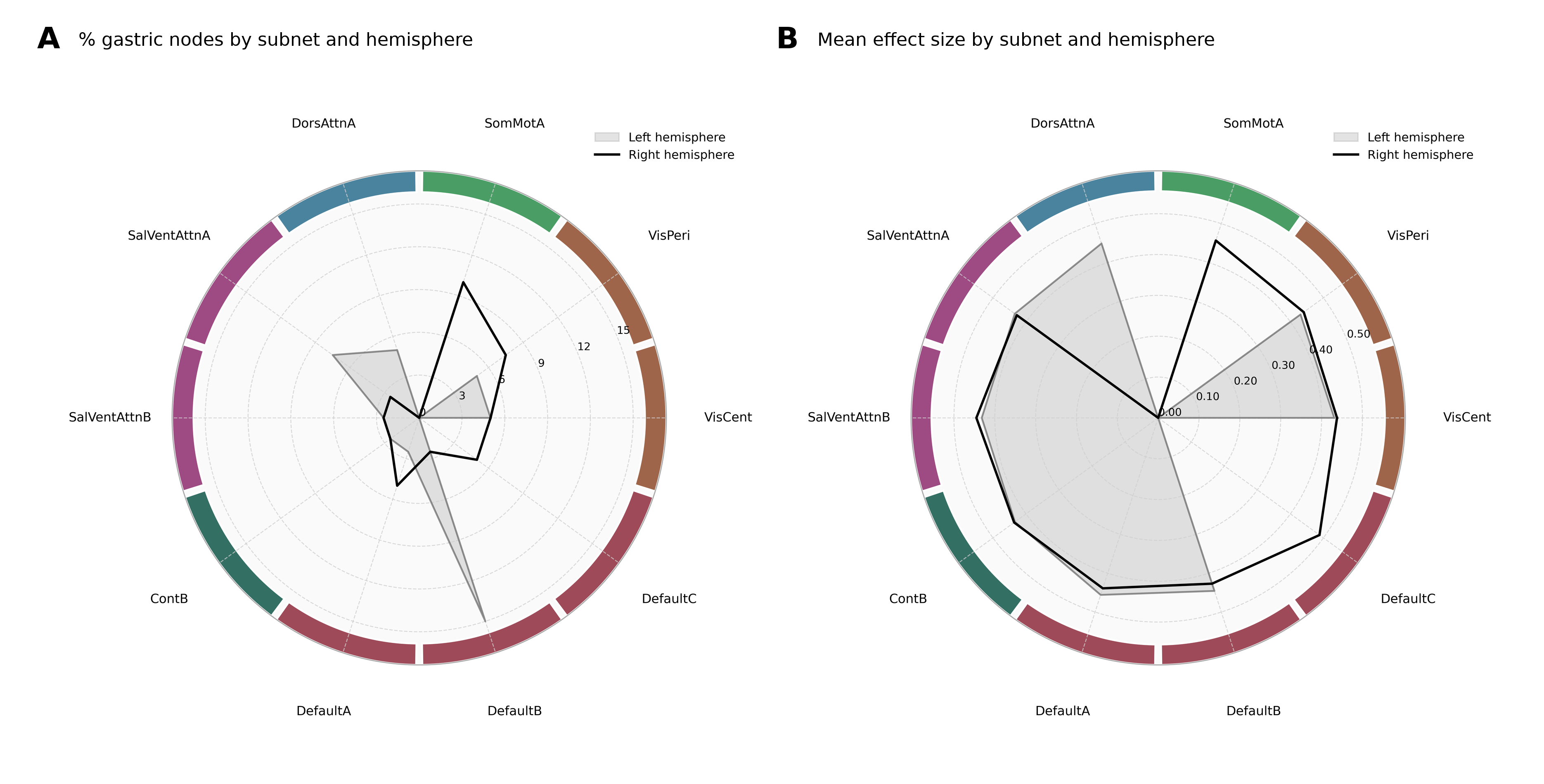

Left: Radar plot showing the average effect size (Cohen’s d) for gastric-BOLD synchronization across cortical subnetworks, plotted separately for the left (grey) and right (black) hemispheres. Shaded areas highlight the relative magnitude and pattern of effects across networks. Numerical scale plotted along each panel denotes levels of effect size from 0 (center point) to maximum value of 0.5. Right: Radar plot depicting the percentage of gastric-responsive nodes within each RSN subnetwork, separated by hemisphere. The plot illustrates how the spatial distribution of gastric-coupled parcels varies across functional systems, revealing both hemispheric asymmetries and network-specific clustering (notably in Default-related and Ventral Attention networks). Numerical scale plotted along each panel denotes percentage values, from 0 (center point) to maximum value of 15%. Note that the plot presents yeo-17 subdivisions of canonical RSN, therefore reflecting lower counts of nodes in each sub-group. Additionally, note that for clearer visualization and finer resolution of network distribution, here we present the yeo-17 RSN network subdivisions, and not the canonical yeo-7 used throughout the paper. Network labels follow the Yeo 17-network cortical parcellation: VisCent and VisPeri denote central and peripheral visual networks; SomMotA-B, somatomotor networks; DorsAttnA-B, dorsal attention networks; SalVentAttnA-B, salience/ventral attention networks; ContA-C, control networks; DefaultA-C, default mode networks; and LimbicA-B, limbic networks.

**Supplementary Figure S2: Temporal circular rotation permutation test for evaluating the statistical significance of burst detection in edge-time series:**

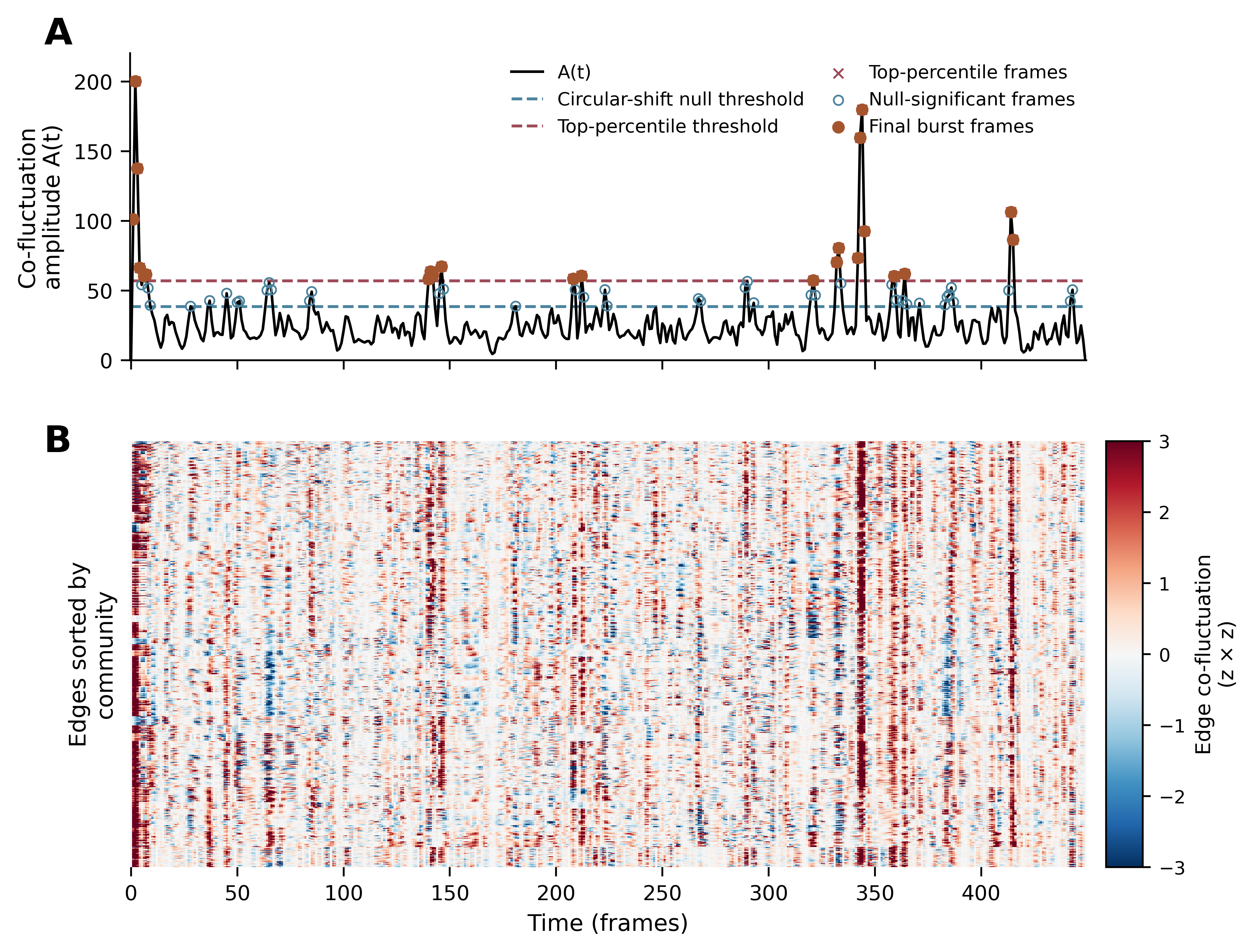

Illustration of the methodology applied for burst detection in one exemplary subject and run (subject AmK, run 2). (A) Observed amplitude A(t) of eTS is presented on the graph as a continuous blue line. To assess burst frames, we used both top 5% percentile and a circular-shift permutation-based thresholds (see *Burst Network Construction* in Methods). The top 5% percentile threshold is marked in fine dashed blue line, while the threshold of the surrogate distribution (null mean = 27.15 ± 6.14) is presented in thick dashed blue line. Event-frames exceeding the top 5% threshold are presented in orange cross, while surrogate significant frames are shown as blue points. Burst frames were defined as points that exceeded both thresholds and are presented on the graph as green points.

(B) After detecting all burst frames, carpet plots of all edges’ eTS were generated and presented throughout the entire scan (450 timeframes), with burst moments represented with darker red in the heatmap, indicating stronger co-fluctuation across multiple edges in the network. Heatmap scale represents edge co-fluctuation intensity in z-score (ranging from -3 to 3).

**Supplementary Figure S3: Thresholding consistency validation at top 10, 15 ,25% amplitude level:**
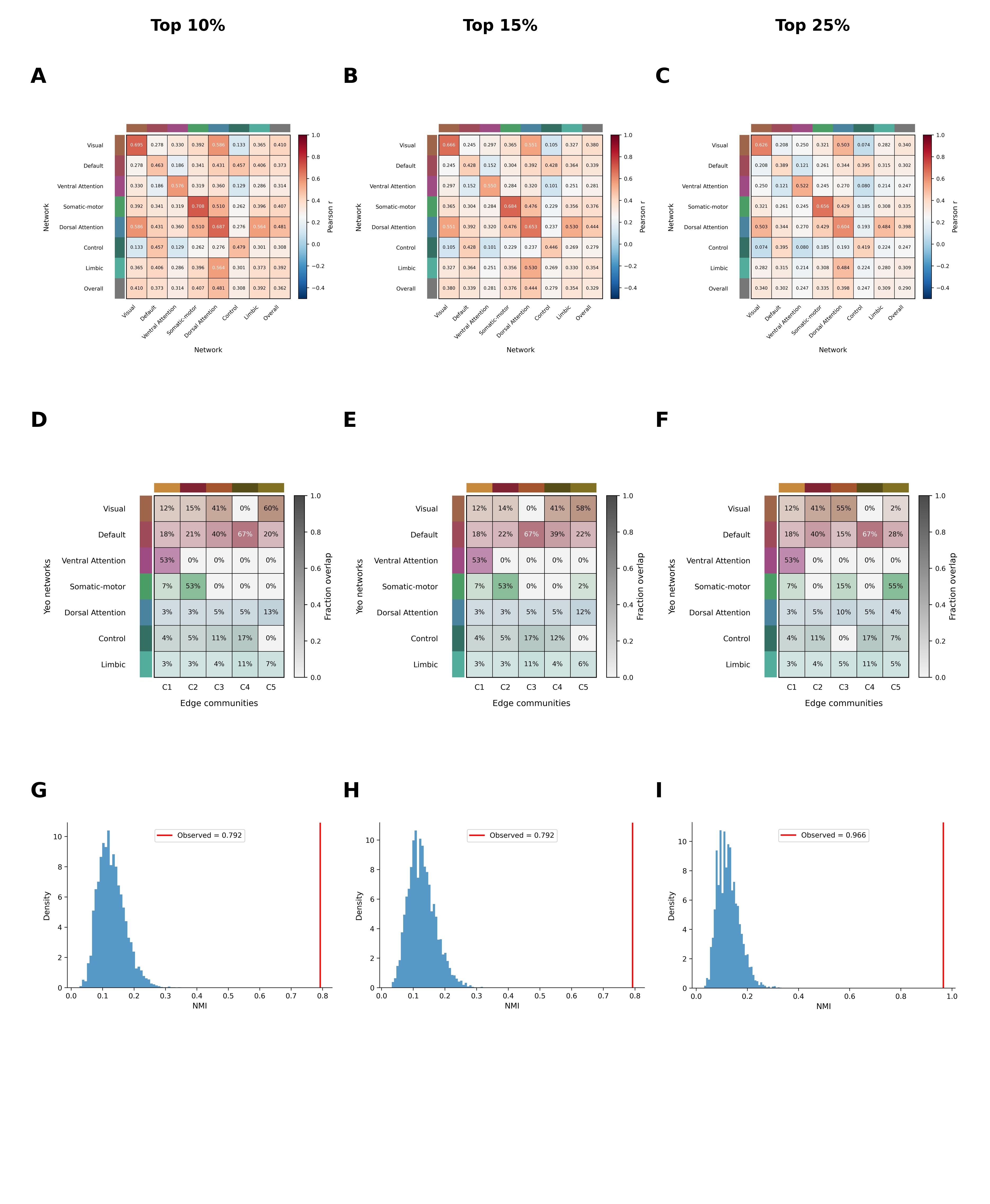

To validate the consistency of results obtained using the primary 5% burst threshold, analyses were repeated using more liberal thresholds (10%, 15%, and 25% of frames based on cofluctuation amplitude). As expected, this resulted in progressively larger numbers of detected burst frames across scan durations (15-minute and 10-minute runs), while preserving the overall structure of the dynamic network.

Top panels show the resulting community structures and their overlap across thresholds. Bottom panels display null distributions of normalized mutual information (NMI) obtained from permutation testing, comparing community partitions derived from the primary 5% threshold with those obtained at higher thresholds.

Across all comparisons, community structure was highly consistent (5% vs. 10%: NMI = 0.792, p = 0.0002, z = 15.50; 5% vs. 15%: NMI = 0.792, p = 0.0002, z = 15.49; 5% vs. 25%: NMI = 0.792, p = 0.0002, z = 15.31). In all cases, observed NMI values (red lines) lay far outside the corresponding null distributions, indicating that the similarity between partitions is significantly greater than expected by chance.

**Supplementary Figure S4: Comparison of observed eFC with phase-randomized surrogate eFC:**

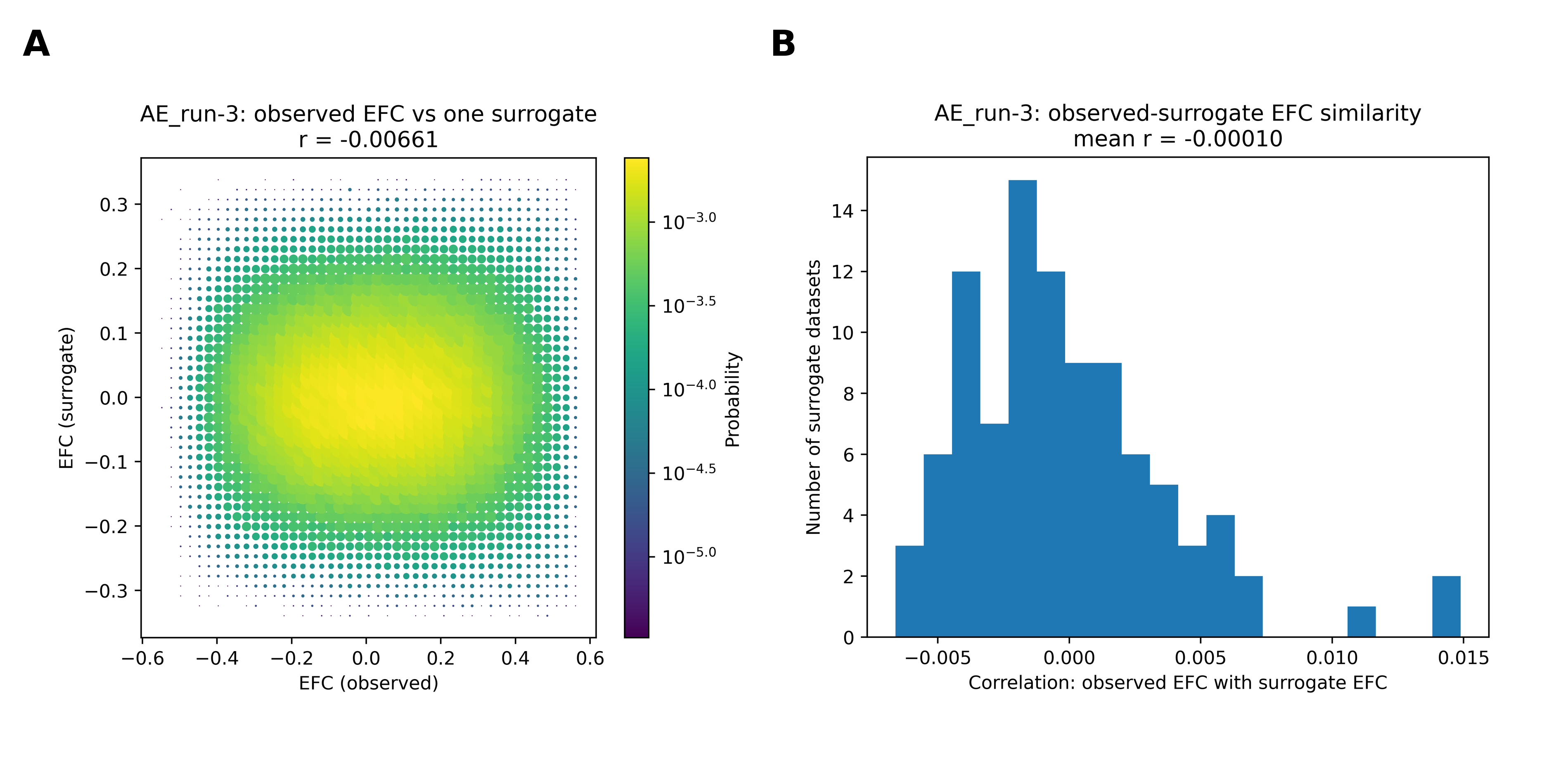

Here, we present the results from the phase-randomization procedure. (A) A two-dimensional histogram from a single subject, comparing observed eFC with eFC for one run from a representative surrogate realization. Point size and color reflect probability density. (B) Distribution of similarity (Pearson correlation) between observed and surrogate eFC across 100 independent realizations. The concentration of probability density around the origin in panel A, together with the distribution of correlation values centered near zero across surrogate datasets (panel B), indicates that observed and surrogate eFC are largely uncorrelated (r = −0.00661; mean r = −0.0001).

**Supplementary Figure S5: Information (NMI) for community-structure comparisons.**

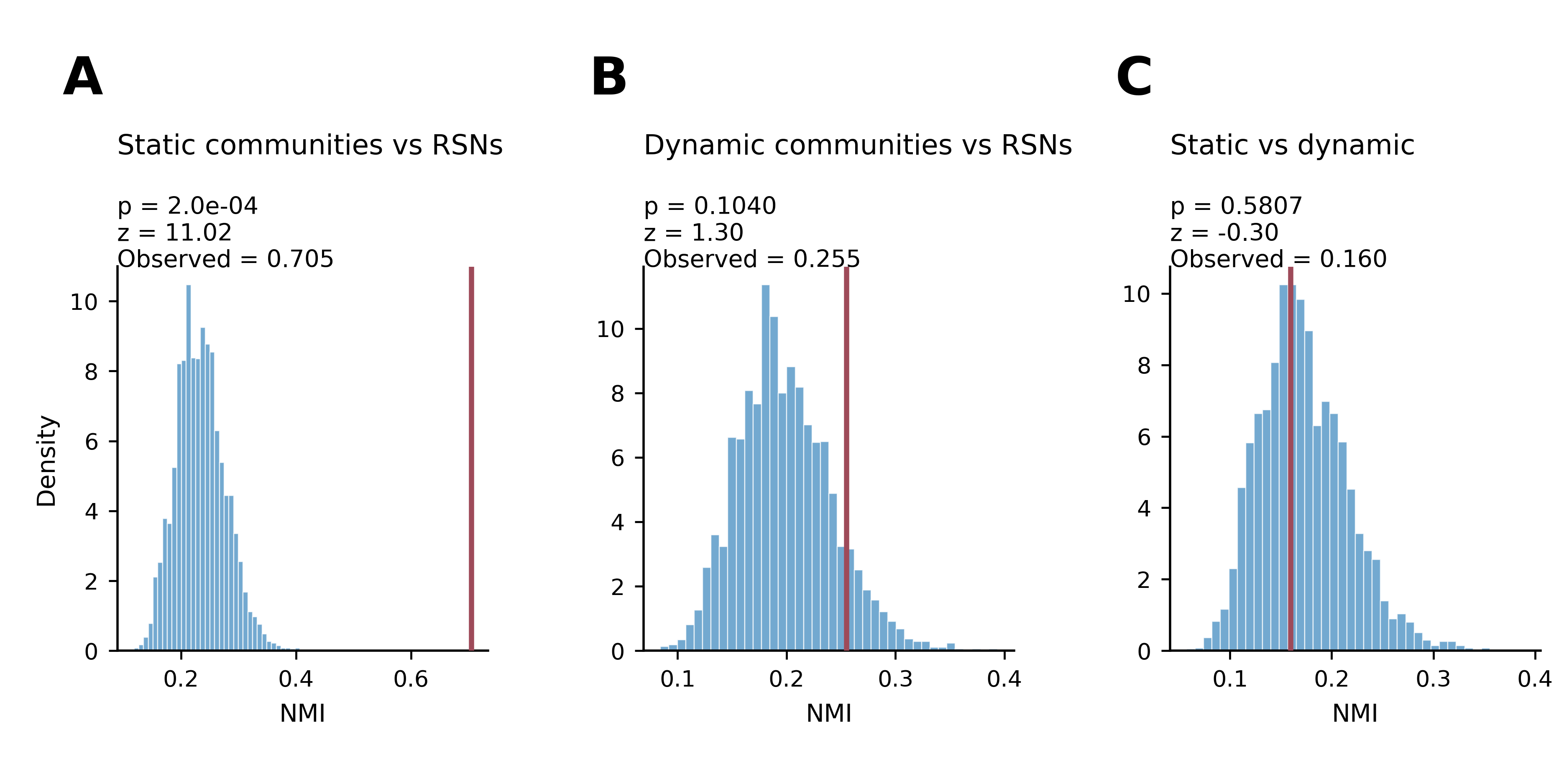

Null distributions were generated using 5,000 node-label permutations to evaluate whether observed community correspondences exceeded chance. The red vertical line marks the observed NMI value in each comparison. (A) NMI between time-averaged gastric-network communities and canonical Yeo-7 resting-state networks. The observed NMI (0.7047) lay far beyond the permutation distribution (p = 0.0002; effect size z = 11.020), indicating a strong correspondence between time-averaged community structure and canonical RSN architecture. (B) NMI between dynamic edge-centric communities and Yeo-7 resting-state networks. The observed NMI (0.255) fell within the null distribution (p = 0.104; z = 1.303), indicating that dynamic communities do not significantly align with canonical RSN modularity. (C) NMI between time-averaged node-based communities and time-varying communities. The observed NMI (0.1595) was not different from chance (p = 0.5807; z = −0.301), demonstrating that time-averaged and dynamic partitions capture distinct organizational regimes
